# 3D Bioprinted Fat-Myocardium Model Unravels the Role of Adipocyte Hypertrophy in Atrial Dysfunction

**DOI:** 10.1101/2025.08.20.671391

**Authors:** Lara Ece Celebi, Pinar Zorlutuna

## Abstract

Cardiovascular diseases (CVD) are the leading cause of mortality in individuals with obesity. Epicardial adipose tissue (EAT) dysfunction serves as a link between obesity and CVD, promoting inflammatory and metabolic alterations that increase CVD risk. While EAT normally supports cardiac health, obesity-induced adipocyte hypertrophy triggers excessive fatty acid and cytokine release, driving myocardial lipotoxicity and inflammation that impair electrophysiology and metabolism, leading to beating irregularities, insulin resistance, and heart failure. The lack of sufficient EAT in small animal models and the impracticality of using large mammals hinder insights into the effects of EAT hypertrophy on the myocardium. To address this gap, a human-derived 3D bioprinted coculture of obese adipocytes and cardiomyocytes (CMs) is developed using patient-derived adipocytes and human induced pluripotent stem cell (hiPSC)-derived atrial CMs (a-iCMs). This platform enables the investigation of both cell-cell and paracrine interactions between hypertrophic adipocytes and a-iCMs, allowing assessment of electrophysiological, structural, and proteomic changes to uncover mechanisms linking EAT hypertrophy to obesity-related atrial dysfunction. Screening of metformin, a cardioprotective drug, reveals improvement in electrophysiological function in hypertrophic adipocyte–a-iCM cultures. 3D bioprinted fat–myocardium model provides a high-throughput platform to study obesity-induced atrial dysfunction and facilitate discovery of therapies for the obese heart.

## 1. Introduction

Obesity has reached pandemic levels by affecting over 1 billion people, equivalent to one in eight worldwide^1^. Cardiovascular diseases (CVD) are the leading comorbidity associated with a high body mass index (BMI) and are responsible for two-thirds of obesity-related deaths^2^. Cardiac adipose tissue dysfunction serves as a link between obesity and CVD, mediating the inflammatory and metabolic alterations that increase CVD risk in obese patients^3^. Unlike other visceral fat depots, epicardial adipose tissue (EAT) lacks distinct boundaries with neighboring tissue^4^ and shares a direct microcirculation with the myocardium. The strategic location of EAT supports cardiac function by facilitating dynamic interactions with the myocardium, including the efficient supply of free fatty acids, responsible for over 60% of the energy required for healthy heart contraction^5^. It also secretes adipokines that regulate endothelial function^6^, exert anti-inflammatory^7^, insulin-sensitizing^6^ and anti-atherosclerotic effects^8^, and contribute to cardioprotection through paracrine and vasocrine signaling mechanisms. However, in obesity, EAT’s strategic location becomes a critical liability, as its expansion (via hyperplasia and hypertrophy) and dysfunction lead to the excessive release of FFAs and proinflammatory mediators directly into the myocardium, disrupting metabolic homeostasis and contributing to lipotoxicity, inflammation, and cardiac dysfunction^9,10^. While healthy adipose tissue expansion favors hyperplasia which is typically accompanied by a proportional angiogenic response, appropriate extracellular matrix (ECM) remodeling, and minimal inflammation^17^, hypertrophic expansion drives obesity-associated metabolic dysfunction^11^. Hypertrophic adipocytes are characterized by several hallmarks: enlarged cell size due to excessive lipid accumulation^12^, increased secretion of proinflammatory cytokines^13^, a shift in adipokine profile^14^, impaired insulin signaling^15^ and altered lipolysis^16^ as well as mitochondrial dysfunction^17^. These changes promote ectopic fat accumulation in the myocardium^18^, metabolic dysfunction and inflammatory signaling within myocardial cells^19^, leading to insulin resistance^20^, arrhythmias^21,22,23,24^, myocardial fibrosis^25,26,27^, and atherosclerosis^28^, culminating in heart failure^29^.

Most clinical studies linking EAT hypertrophy to CVD have utilized imaging techniques such as computed tomography and magnetic resonance imaging^30^. While useful for associating risk factors, these imaging studies often lack mechanistic insights into disease progression since they are primarily retrospective^31^. To bridge this gap, several mechanistic explanations have been proposed utilizing *in vitro* and *in vivo* platforms. Venteclef et al. investigated the paracrine effects of human EAT on rat cardiomyocytes (CMs) utilizing organo-culture model of rat atria and demonstrated that adipo-fibrokines released by EAT can promote myocardial fibrosis^25^. This finding suggests that hypertrophic EAT may actively contribute to adverse cardiac remodeling by modulating the local myocardial environment through paracrine signaling. However, small lab animals have limitations in modeling the contractile function of the human heart due to their small size and short lifespan^32^. More importantly, there are notable differences in cardiac fat depots, as most commonly used small lab animals, mice and rats, possess minimal EAT, which is primarily restricted to the atrioventricular groove, and with no direct contact with the myocardium^33^, which limits their translational relevance. In contrast, large animal models such as pigs and sheep more closely mimic human EAT distribution and cardiac physiology. Despite their physiological relevance, large animal models are low-throughput and costly, limiting their utility for mechanistic studies and early-stage therapeutic screening^34^. Therefore, scalable human-derived *in vitro* systems are critical to complement and inform animal studies in the search for effective therapeutics^35^.

Human induced pluripotent stem cells (hiPSCs) provide physiologically relevant platforms that potentially enable mechanical investigations, personalized approaches for drug testing, and cytotoxicity assessments. Recently, researchers cocultured adipose-derived stromal cells (ADSC)-derived human adipocytes with hiPSC-derived ventricular CMs (v-iCMs) in 2D settings^36^. They observed prolonged action potential duration, extended calcium transients, reduced conduction velocity, and increased conduction velocity heterogeneity, along with altered levels of ion channel and gap junction genes in 2D v-iCMs co-cultured with adipocytes or exposed to adipocyte-conditioned medium. This study also demonstrated that paracrine signals from adipocytes alone are sufficient to induce electrophysiological dysfunction. However, 2D cultures of adipocytes and CMs have some inherent limitations. In 2D, adipocytes often lose their natural spherical shape, which can affect gene expression, lipid storage, and adipokine secretion^37^. CMs in monolayer lack the maturation seen in native cardiac tissue, limiting their ability to accurately replicate physiological contraction and electrophysiology^38^. Additionally, without the ECM architecture found *in vivo*, 2D systems may not fully capture the complex cell-cell and cell-matrix interactions involved in adipose tissue dysfunction in obesity. Moreover, the adipocytes used in existing monolayer coculture models with CMs^36,39,40^ have yet to recapitulate the dysfunctional phenotype characteristic of obesity.

Chamber specificity is crucial for modeling cardiac pathologies, as each heart chamber has distinct structural, functional, and electrophysiological characteristics that influence the manifestation and progression of CVD^41^. However, most hiPSC-derived engineered models have predominantly focused on generating ventricular myocardium, overlooking the need for atrial myocardium models^42^. Obesity is associated with a 50% increase in the risk of developing atrial fibrillation^43^. Therefore, there is an urgent need for engineered tissues that accurately replicate atrial dysfunction in obesity, providing chamber-specific mechanistic insights and facilitating the development of targeted therapeutic strategies. Here, we present a human-derived 3D adipocyte-hiPSC derived atrial CM (a-iCM) model designed to enable precise quantification of both contact-dependent (juxtacrine) and paracrine interactions within the context of obesity, which is crucial for the development of targeted and effective therapeutic interventions aimed at treating obesity-related heart failure. As reported recently^35^, there is no existing literature on 3D coculture systems combining adipocytes and CMs. Therefore, this represents the first 3D coculture system combining adipocytes and CMs, established using human patient-derived cells and high-fat diet (HFD) to recapitulate the obese adipocyte phenotype to uncover mechanisms linking EAT dysfunction to atrial impairment in obesity.

We previously employed extrusion-based 3D bioprinting and gelatin methacryloyl (GelMA)-Collagen type I bioink to model (patho)physiological characteristics of ventricular myocardium^44^. Here, we once again utilized 3D bioprinting and the characterized GelMA-Collagen type 1 bioink^44^ to engineer 3D cultures of healthy (Adi) and high-fat diet (HFD)-induced hypertrophic (obese-like, OAdi) ADSC-derived adipocytes. We characterized the major hallmarks of hypertrophic adipocytes and their paracrine effects on mitochondrial and glycolytic function as well as ATP production in a-iCMs. Next, we 3D bioprinted a-iCMs within Adi and OAdi constructs, creating a 3D coculture. Using this coculture model, we reported the effects of 3D Adi/OAdi on a-iCMs, examining changes in electrophysiology, morphology, and protein expressions related to atrial dysfunction in obesity.

Metformin is an antidiabetic drug that lowers blood glucose levels by activating AMP-activated protein kinase (AMPK), which suppresses hepatic gluconeogenesis, enhances insulin sensitivity, and promotes glucose uptake in peripheral tissues^45^. To demonstrate the utility of the model for therapeutic screening, we evaluated metformin’s ability to mitigate dysfunction in a-iCMs exposed to hypertrophic adipocytes. In the heart, metformin’s activation of AMPK improves cardiac function by reducing inflammation and oxidative stress^46^, enhancing fatty acid oxidation^47^, protecting against insulin resistance^48^, and mitigating fibrosis^49^. At the CM level, low-dose metformin (<2.5 mM) enhances cellular respiration by stimulating mitochondrial biogenesis via AMPK signaling^50^. It also preserves mitochondrial membrane potential^51^, reduces reactive oxygen species (ROS) production^51^, and regulates autophagy^52^ to reverse obesity-related CM dysfunction. At the adipocyte level, metformin reduces circulating FFAs and adipose tissue lipolysis^53^. These effects collectively protect CMs from lipotoxicity, metabolic stress, and contractile dysfunction associated with obesity-induced cardiac injury. Using 3D bioprinted constructs, we tested whether metformin treatment could mitigate hypertrophic adipocyte-induced electrophysiological and metabolic impairments in a-iCMs.

In this study, we established a human-derived 3D bioprinted model that integrates ADSC-derived hypertrophic adipocytes and a-iCMs to replicate key features of obesity-related cardiac dysfunction. We demonstrated that hypertrophic adipocytes disrupt a-iCM metabolic function, contractility, calcium handling, and insulin signaling through both paracrine and/or juxtacrine interactions. Furthermore, we validated the therapeutic screening potential of constructs by demonstrating metformin’s ability to modulate electrophysiological and metabolic impairments in 3D OAdi-a-iCMs. Engineered EAT-myocardium models provide a biomimetic platform to investigate the mechanisms underlying obesity-induced cardiac dysfunction and enable high-throughput drug discovery to develop targeted treatments for CVD comorbidities.

## 2. Results and Discussion

### 2.1. 3D Hypertrophic Adipocytes Display Hallmarks of Obesity

#### 2.1.1. Lipid Droplet Enlargement and Cytoskeletal Changes

*In vivo*, obese adipocytes are characterized by an increased capacity for triglyceride storage, leading to fewer but larger lipid droplets^54^, which is indicative of a unilocular morphology typical of mature white adipocytes^55^. Previously, this phenomenon was replicated *in vitro* by exposing adipocytes to a fatty acid-rich environment^16^, which induces features similar to an *in vivo* obese phenotype. Here, we 3D bioprinted and differentiated ADSCs into adipocytes representing healthy and obese states by mimicking a high-fat diet (HFD) through palmitic acid (PA) supplementation to induce adipocyte hypertrophy (Figure 1A). To assess lipid accumulation, we performed Bright field (BF) imaging and Nile Red staining after a 6-day PA treatment, revealing an increase in lipid droplet size in the obese constructs (3D OAdi) compared to controls (3D Adi). BF images demonstrated clear morphological distinctions, with 3D OAdi exhibiting larger lipid droplets compared to the control (Figure 1B, SFigure 1B). Nile Red staining confirmed the accumulation of lipids, with 3D OAdi showing larger lipid droplet formation compared to the control (Figure 1C, SFigure 1C-D).

**Figure 1:**
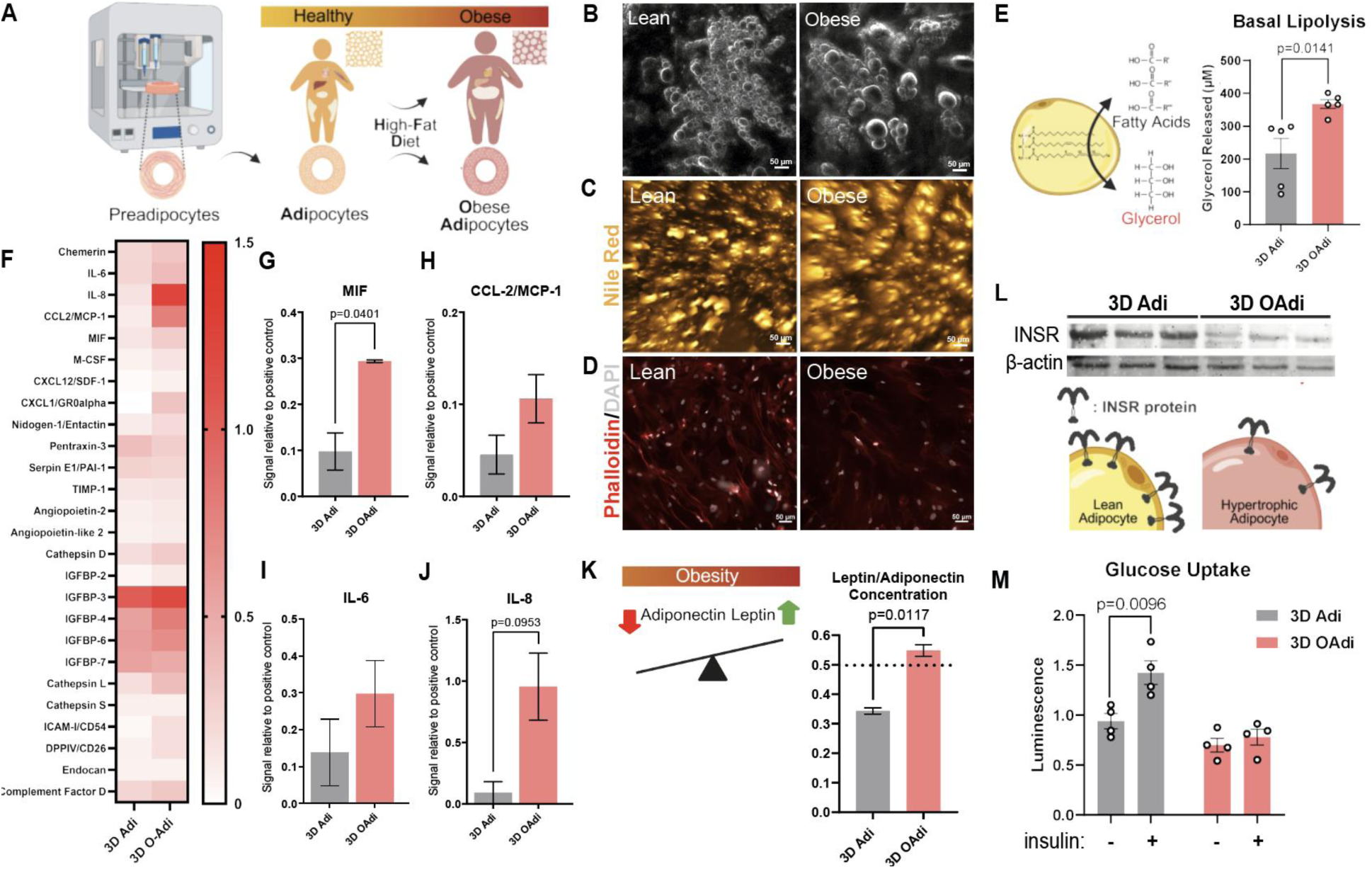
Characterization of 3D Adi and OAdi constructs according to well-established hallmarks of obese adipocytes. A) Schematic displaying hypertrophy progression of obese adipocytes and recapitulation of this process using engineered constructs. B) Bright-field imaging *(scale bar=50 um)*, C) Nile Red staining *(scale bar=50 um),* and D) F-actin staining *(scale bar=50 um)* of 3D Adi/OAdi. E) Glycerol release per 3D Adi/OAdi construct over 48 hours (n=5). F) Human cytokine array: G) MIF (n=2), H) MCP-1 (n=2), I) IL-6 (n=2), J) IL-8 (n=2), and K) Leptin/Adiponectin ratio (n=2) quantification in 3D Adi/OAdi secretome. L) INSR expression (n=3) and M) Basal and insulin-dependent glucose uptake (n=4) of the 3D Adi/OAdi.

The organization of F-actin around the 3D Adi and OAdi constructs was evaluated by F-actin staining, revealing altered cortical actin structures in the obese constructs relative to the control, recapitulating the cytoskeletal changes driven by obesity-induced ECM remodeling (Figure 1D). Cytoskeletal alterations observed in PA-treated adipocytes may be due to the adaptive response to accommodate lipid droplet enlargement in hypertrophic adipocytes. The alteration of actin cytoskeleton organization, particularly the loss of cortical F-actin due to lipid droplet unilocularization, was reported to compromise adipocyte’s ability to respond to insulin and effectively translocate glucose transporters, thereby impairing glucose uptake and contributing to the development of insulin resistance^56^. These results confirm that lipid droplet enlargement in 3D OAdi is accompanied by cytoskeletal alterations, supporting key structural hallmarks expected in obesity-associated adipocyte dysfunction.

#### 2.1.2. Secretome Analysis and Cytokine Profiling

Lipolysis is the metabolic process by which triglycerides, stored in adipocytes, are broken down into FFA and glycerol. In a healthy heart, FFA secreted by cardiac fat depots supply over 60% of the energy needed for contractile function^5^. In lean adipocytes, the processes of energy uptake and release are tightly regulated, maintaining balanced lipolysis, where triglycerides are hydrolyzed into FFAs and glycerol as needed^57^. In contrast, hypertrophic adipocytes exhibit elevated basal lipolysis, resulting in an increased breakdown of triglycerides and FFA release^58^. When FFAs are chronically elevated, excess fatty acids accumulate in neighboring tissues, including the myocardium, which lacks the fat storage capacity to handle them efficiently^59^. We evaluated the basal lipolytic activity of the 3D Adi/OAdi by measuring glycerol release from the constructs as an indicator of lipolysis. 3D OAdi exhibited significantly increased basal lipolysis compared to 3D Adi (p=0.0141, Figure 1E), as measured by elevated glycerol release. This finding aligns with *in vivo* dyslipidemic profile^60^ and supports *in vitro* findings on 3D HFD-induced hypertrophic ADSC-derived adipocytes^16^.

Macrophage migration inhibitory factor (MIF), IL-6, and IL-8 are proinflammatory cytokines that play a key role in modulating immune responses, promoting macrophage retention, and contributing to chronic inflammation, which are all hallmarks of obesity-related tissue dysfunction^61,62,63^. Human cytokine array analysis (Figure 1F) revealed significantly higher levels of MIF (Figure 1G, p=0.0401), substantially increased concentrations of other proinflammatory cytokines such as IL-6 (Figure 1I) and IL-8 (Figure 1J), as well as cytokines associated with ECM remodeling such as MCP-1 (Figure 1H), in 3D OAdi compared to the lean control.

Leptin and adiponectin are adipokine hormones that play key roles in metabolic regulation^64^. Leptin normally suppresses appetite and promotes energy expenditure^65^, while adiponectin enhances insulin sensitivity and has anti-inflammatory properties^6^. Another consequence of adipocyte hypertrophy in obesity is the decrease in adiponectin secretion and a corresponding increase in leptin release, disrupting metabolic homeostasis and promoting insulin resistance and inflammation^66^. Under physiological conditions, the leptin-to-adiponectin ratio is typically below 0.5, and values exceeding this threshold have been associated with an increased risk of CVD^67^. 3D OAdi constructs had a higher leptin to adiponectin concentration ratio compared to the lean control and exceeded the 0.5 threshold (0.53 ± 0.04, p=0.0117, Figure 1K). Taken together, these findings indicate that 3D OAdi constructs effectively recapitulate obesity-induced alterations in the adipose tissue secretome, including enhanced lipolysis, inflammatory cytokine secretion, and hormonal imbalance.

#### 2.1.3. Insulin Receptor Prevalence and Activation

Adipocytes express insulin receptors (INSR) that mediate glucose uptake, lipid metabolism, and energy storage in response to insulin signaling^68^. Obesity is the most common cause of insulin resistance, a condition in which insulin-sensitive tissues, such as adipose tissue, skeletal muscle, and liver, fail to respond adequately to circulating insulin^69^. In individuals with obesity or diabetes, this impaired response is attributed to reduced surface expression of the insulin receptor (INSR) and diminished INSR kinase activity^70^, leading to the development of type 2 diabetes mellitus (T2D) and CVD. INSR expression is decreased in the adipose tissue of obese patients compared to normal-weight controls, with a progressive decline observed as BMI increases^69^. Similar to *in vivo* results, HFD-induced 3D OAdi had reduced INSR expression compared to the lean control (Figure 1L). Notably, this reduction in INSR expression was observed in the absence of systemic inflammation or immune cell involvement, indicating that fatty acid-induced adipocyte hypertrophy alone is sufficient to impair insulin receptor expression in adipocytes.

To determine whether reduced INSR expression in 3D OAdi translates to impaired metabolic function, we assessed insulin-stimulated glucose uptake as a functional readout of insulin responsiveness. While insulin treatment significantly increased glucose uptake in 3D Adi constructs (p = 0.0096), 3D OAdi showed no significant response, indicating loss of insulin sensitivity in hypertrophic adipocytes (Figure 1M). Additionally, there was a substantial decrease in basal glucose uptake in 3D OAdi constructs compared to 3D Adi controls (p=0.0577, Figure 1M). This could be an outcome of increased reliance on fatty acid oxidation due to HFD, or alternatively, a result of impaired basal glucose transporter expression and function.

Combined with elevated basal lipolysis, these findings could point to the presence of a vicious cycle, where elevated FFAs impair insulin signaling, and insulin resistance further promotes lipolysis^60^. Additionally, the increased leptin-to-adiponectin ratio, which has been proposed as a marker of insulin resistance^71^, may be a contributor to the disrupted glucose metabolism and insulin insensitivity we observed in 3D OAdi. Our findings are consistent with prior *in vitro* (murine adipocyte) and *in vivo* studies using fatty acid-treated adipocytes and short-term HFD-fed mice^15^, which demonstrated that adipocyte hypertrophy alone, induced without immune cell infiltration or systemic inflammation, is sufficient to impair insulin receptor expression and disrupt insulin-stimulated glucose uptake. These findings indicate that 3D OAdi constructs effectively model obesity-induced changes in adipose tissue, including enhanced lipid storage, structural remodeling, increased proinflammatory signaling, disrupted insulin signaling, and insulin-dependent glucose uptake, making them a biomimetic platform for studying obesity-induced cardiac dysfunction.

### 2.2. 3D Hypertrophic Adipocyte Secretome Induces Metabolic Dysfunction in Atrial Cardiomyocytes

#### 2.2.1. Mitochondrial Bioenergetic Function

Obese patients have alterations in myocardial substrate utilization^72^ and reduced ATP production/utilization efficiency^73^, indicating mitochondrial dysfunction. Earlier studies suggested that CM mitochondrial dysfunction is caused by myocardial lipid accumulation, impaired glucose tolerance, and insulin resistance^74^. Here, we evaluated and compared the paracrine effects of 3D Adi and 3D OAdi on the mitochondrial function of a-iCMs using Seahorse extracellular flux analysis (Figure 2A). We observed that 3D OAdi conditioned media (ConM) treated a-iCMs exhibited significantly reduced basal respiration compared to a-iCMs cultured with either the B22 (control) (p=0.0111) or 3D Adi secretome (p=0.0005, Figure 2D). Moreover, a-iCMs cultured with the 3D OAdi ConM demonstrated a significant reduction in maximal respiration (p=0.0320, Figure 2E) and spare respiratory capacity (p=0.0044, Figure 2F) compared to 3D Adi group, which indicates a diminished ability of the mitochondria to adapt to increased energy demands during periods of stress. Coupling efficiency, which reflects how effectively mitochondria convert electron transport chain activity into ATP production, was also significantly reduced in a-iCMs cultured with the 3D OAdi ConM, both compared to the B22 control and 3D Adi group (p<0.0001, Figure 2G). Furthermore, ATP-production coupled respiration was significantly decreased in a-iCMs cultured with the 3D OAdi ConM, both compared to the B22 control (p=0.0014) and 3D Adi group (p=0.0002, Figure 2H). Mitochondrial ATP production (mitoATP), reflecting the ATP generated through mitochondrial oxidative phosphorylation (OXPHOS), was significantly lower in 3D OAdi ConM-treated a-iCMs compared to 3D Adi (p=0.0186, Figure 2I). Importantly, mitoATP of a-iCM treated with 3D Adi was significantly increased compared to B22 control, indicating secreted factors from healthy adipocytes may enhance mitochondrial OXPHOS and support cardiomyocyte energy metabolism. Taken together, these findings suggest that the secretome of hypertrophic adipocytes impairs mitochondrial function of a-iCMs through paracrine mechanisms, evidenced by reduced basal and maximal respiration, spare capacity, coupling efficiency, and mitochondrial ATP production. In contrast, secreted factors from lean adipocytes promote a metabolic shift from glycolysis toward OXPHOS, potentially priming a-iCMs toward a more mature energetic phenotype seen *in vivo*^75^.

**Figure 2:**
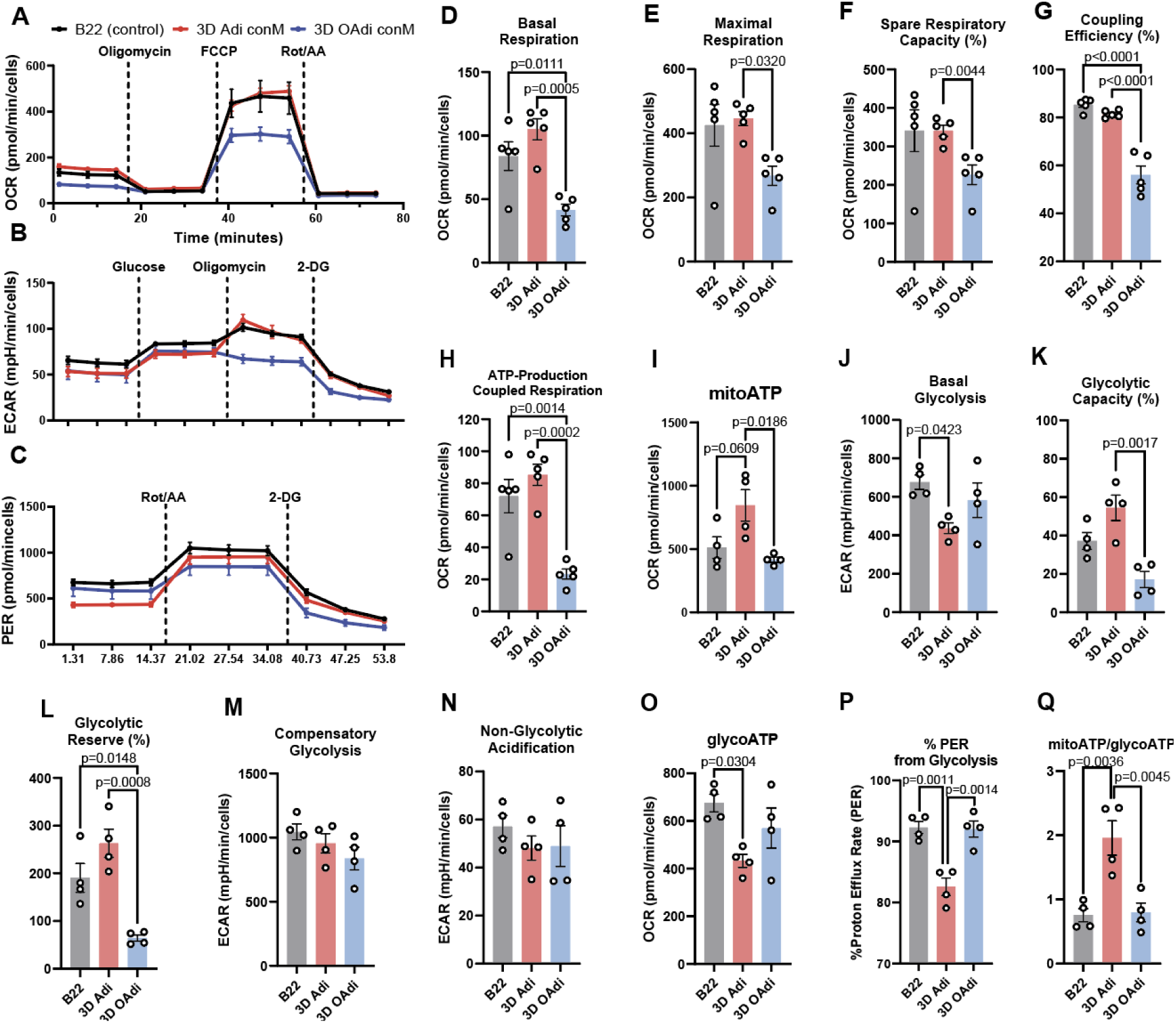
Metabolic profiling of a-iCMs exposed to 3D Adi/OAdi ConM reveals impaired mitochondrial function and altered glycolytic activity. A) Oxygen consumption rate (OCR) following sequential addition of oligomycin, FCCP, and rot/AA (Seahorse Mito Stress test) (n=5). B) Extracellular acidification rate (ECAR) following glucose, oligomycin, and 2-DG injections (Seahorse Glycolysis Stress test) (n=4). C) Proton efflux rate (PER) analysis rot/AA, and 2-DG injections (Seahorse Glycolytic Rate test) (n=4). Quantification of D) basal respiration (n=5), E) maximal respiration (n=5), F) spare respiratory capacity (n=5), G) coupling efficacy, H) ATP-linked respiration (n=5), I) mitochondrial ATP production (mitoATP) (n=4) J) basal glycolysis (n=4), K) glycolytic capacity (n=4), L) glycolytic reserve (n=4), M) compensatory glycolysis (n=4), N) non-glycolytic acidification (n=4), O) glycolysis-dependent ATP production (glycoATP) (n=4), and P) percent PER from glycolysis (n=4) Q) The mitoATP/glycoATP ratio of ConM treated a-iCMs (n=4).

#### 2.2.2. Glycolytic Function

When CMs sustain mitochondrial damage, glycolysis is expected to increase as the impaired mitochondria are unable to generate adequate ATP through OXPHOS^76^. This metabolic flexibility forces the cell to rely on the less efficient but faster glycolytic pathway to meet its energy demands and sustain cellular functions^76^. However, obesity-related cardiometabolic diseases are characterized by metabolic inflexibility^77^, where cells are unable to switch between fuel sources such as glucose and fatty acids. Here, we compare the paracrine effects of 3D Adi and OAdi on the glycolytic function of a-iCMs using Seahorse extracellular flux analysis (Figure 2B-C). The basal glycolysis of 3D Adi-ConM treated a-iCMs was significantly lower compared to the B22 control (Figure 2J, p=0.0423). Combined with the Mito Stress assay results (Figure 2A-I), the decrease in basal glycolysis might reflect an early metabolic shift from glycolysis toward OXPHOS induced by lean adipocyte secreted factors. Glycolytic capacity, which measures the maximum glycolytic potential, was significantly reduced in the 3D OAdi ConM-treated group compared to 3D Adi ConM-treated group (Figure 2K, p=0.0017). Glycolytic reserve, representing the additional glycolytic capacity available under stress, was significantly impaired in 3D OAdi ConM-treated a-iCMs compared to B22 control (p=0.0148) and 3D Adi ConM-treated group (p=0.0008, Figure 2L). Notably, there was a substantial increase in glycolytic capacity and reserve in a-iCMs treated with 3D Adi ConM compared to B22 control (Figure 2K-L), suggesting that while mitochondrial ATP production is enhanced, the capacity for glycolytic compensation is also preserved, and even markedly increased, indicating improved metabolic flexibility.

There was a marked decrease in compensatory glycolysis, which shows reduced ability of CM to shift toward ATP production from glycolysis (glycoATP) when mitochondrial function is impaired in the OAdi group compared to B22 control (Figure 2M). Non-glycolytic acidification, which reflects proton production from sources other than glycolysis, remained similar across the groups (Figure 2N). This suggests that the observed differences in extracellular acidification were primarily driven by changes in glycolytic activity, rather than alterations in fatty acid oxidation or other non-glycolytic metabolic pathways. glycoATP was significantly lower in the 3D OAdi CM-treated a-iCMs compared to the other groups (Figure 2O, p=0.0019). The percentage of proton efflux rate (PER) derived from glycolysis was significantly reduced in the 3D Adi ConM group compared to the B22 control (p=0.0011) and 3D OAdi group (Figure 2P, p=0.0014), which further indicated a diminished reliance on glycolysis for extracellular acidification in 3D Adi group. The ratio of mitoATP to glycoATP production was significantly higher in the 3D Adi group compared to the B22 control (p=0.0036) and 3D OAdi (Figure 2Q, p=0.0045), confirming a metabolic shift toward mitochondrial oxidative phosphorylation induced by lean adipocyte secretome. Taken together, 3D OAdi ConM-treated a-iCMs exhibit marked metabolic inflexibility, characterized by an impaired ability to upregulate glycolysis in response to mitochondrial dysfunction, suggesting that paracrine signals from hypertrophic adipocytes compromise their capacity to adapt to energetic stress. In contrast, 3D Adi ConM-treated a-iCMs displayed enhanced metabolic flexibility, characterized by increased mitochondrial ATP production alongside preserved and elevated glycolytic reserve, suggesting that lean adipocyte secretome supports a balanced and adaptive energy metabolism in a-iCMs.

In our supplementary studies, we also included an expanded analysis with additional control groups, such as two different concentrations of adiponectin and leptin correlating the 3D Adi (adiponectin-high, leptin-low) and OAdi secretome content (adiponectin-low/leptin-high, as well as two different concentrations of PA to reflect the dose-dependent metabolic effects of lipotoxicity. 3D OAdi ConM and Leptin-high treated a-iCMs had significantly lower spare respiratory capacity compared to 3D Adi ConM, indicating the leptin content of 3D OAdi ConM might be contributing to a reduced ability to meet increased energy demands under stress (SFigure 2D). a-iCMs treated with increasing concentrations of leptin exhibited increased glycoATP observed at the highest leptin dose compared to lower concentrations (SFigure 2F), consistent with *in vivo* studies showing leptin enhances glucose utilization^78^. Coupling efficacy and mitoATP production of 3D OAdi ConM treated a-iCMs were significantly reduced (SFigure 2E, G) compared to every other group, which suggests that the changes seen in mitochondrial function of 3D OAdi ConM treated a-iCMs might be due to some other small molecule or a combinatorial effect of the contents of this secretome.

Although glycolysis was relatively similar across groups (SFigure 2I), PA-treated groups exhibited a similar reduction in glycolytic capacity and reserve (%) as observed in a-iCMs exposed to 3D OAdi ConM (SFigure 2J-K), suggesting that the elevated fat content in the 3D OAdi secretome may contribute to impaired glycolytic function in a-iCMs. Lastly, non-glycolytic acidification, typically linked to other cellular or metabolic processes that generate protons, was significantly elevated in the PA-high group, potentially reflecting increased fatty acid oxidation (SFigure 1L). Together, these findings demonstrate that the secretome of hypertrophic adipocytes induces profound metabolic dysfunction in a-iCM by impairing both mitochondrial OXPHOS and glycolytic adaptability, whereas the lean adipocyte secretome promotes metabolic flexibility and a shift toward a more efficient, OXPHOS-dominant energetic phenotype of a-iCMs similar to healthy adult human CM^79^.

Despite the limitations of 2D adipocyte cultures (e.g., detachment from the tissue culture plate during cultivation periods longer than 28 days^80^, dedifferentiation after only one week of culture^81^), we also ran supplementary experiments (SFigure 3-4) where we collected ConM from both 2D adipocytes (2D Adi) and hypertrophic adipocytes (2D OAdi) and applied it to a-iCMs. We observed decreased mitochondrial function in both groups (SFigure 4A-D), with the hypertrophic adipocyte group showing even lower glycoATP levels (SFigure 4E). Overall, culturing adipocytes as monolayers (2D Adi/OAdi) did not yield a clear distinction in their paracrine effects on a-iCMs. We suspect this may be due to adipocytes, even lean ones, adopting a pathological phenotype when maintained on plastic surfaces, thereby impairing both mitochondrial and glycolytic function of a-iCMs, which warrants further investigation.

**Figure 3:**
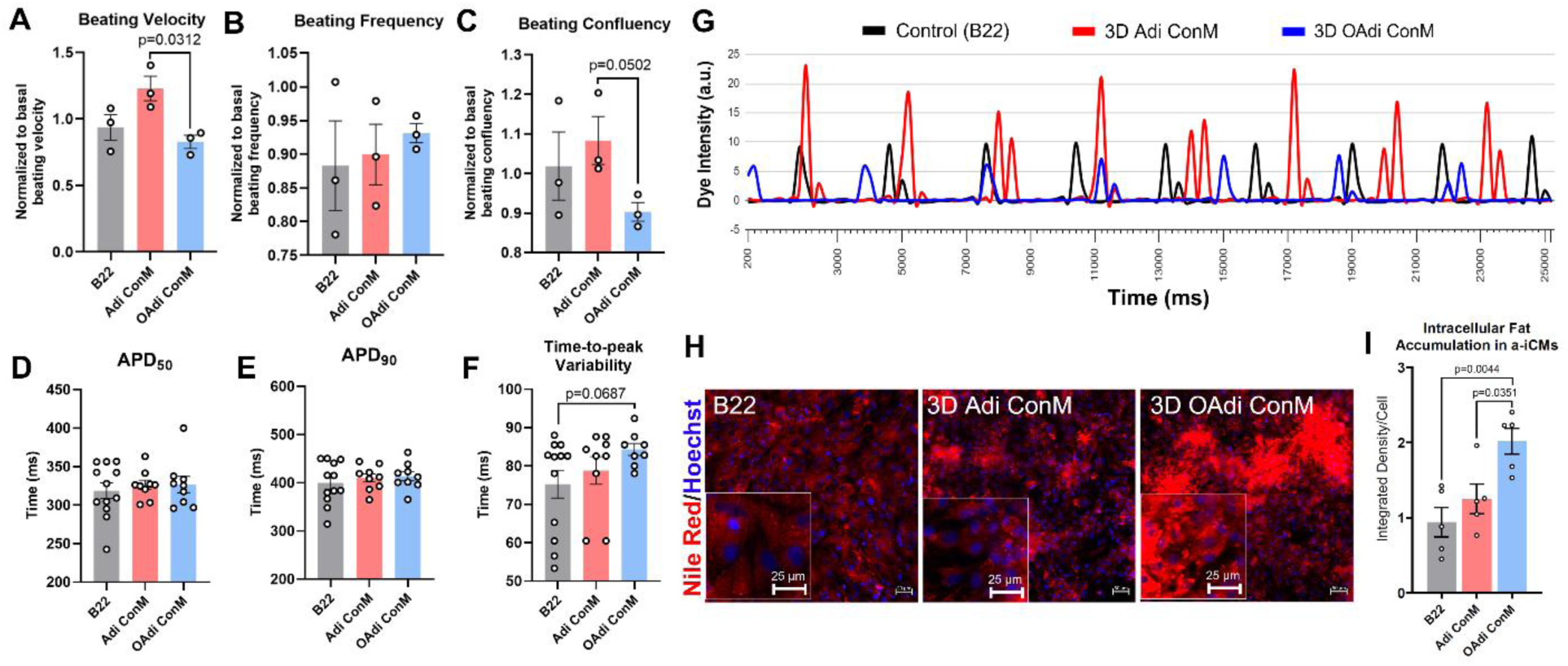
Alteration of beating characteristics in a-iCMs exposed to 3D OAdi conditioned media (ConM) and ectopic fat accumulation. Brightfield beating analysis of A) beating velocity (n=3), B) beating frequency (n=3), and C) beating confluency of a-iCMs treated with ConM (n=3), D) APD50 (n=9-12), E), APD90 (n=9-12), F) time-to-peak variability in a-iCMs treated with ConM (n=8-12), G) representative calcium transient waveforms of B22, 3D Adi/OAdi ConM treated a-iCMs H) Nile Red staining (red: Nile Red, lipid, blue: Hoechst33342, nuclei) *(scale bar=50 um)* and magnified image *(scale bar=25 um)* and I) quantification of integrated density of ectopic lipid accumulation/cell in a-iCMs treated with ConM (n=5).

**Figure 4.**
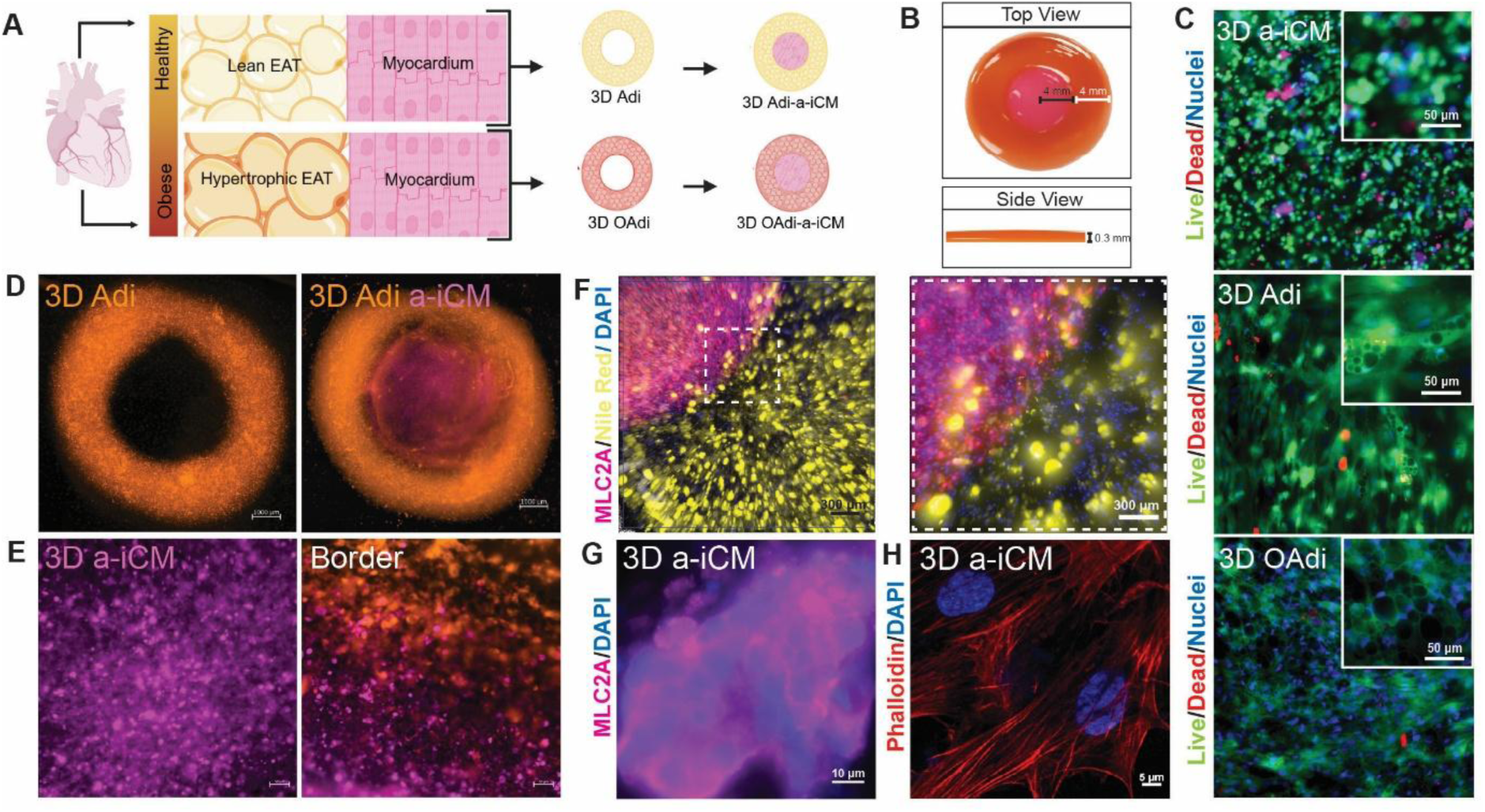
3D bioprinting of Adi/OAdi a-iCM constructs. A) Schematic representation of lean and hypertrophic EAT and the process of 3D bioprinting of Adi/OAdi a-iCM constructs. B) Image of the 3D bioprinted concentric ring construct (8 mm outer diameter, 4 mm inner diameter, 0.3 mm height). C) Live-Dead imaging of 3D construct regions (Green: live cells, red: dead cells, blue: nuclei (hoechst 33342), *Scale bar: 50 μm*). D) Cell colocalization in the constructs using CellTracker (a-iCM: Far Red, Adipocyte: Orange, *Scale bar: 1000 μm*) E) a-iCM and border regions of the construct *(Scale bars: 50 μm)* F) Immunofluorescence images of a-iCM marker (MLC2A, magenta) and adipocyte marker (Nile Red, yellow) *(Scale bar: 300 μm, Magnified Image Scale Bar: 100 μm)* G) MLC2A/DAPI staining confirming atrial iCM identity *(Scale bar: 10 μm)*. H) Phalloidin/DAPI staining visualizing actin cytoskeleton organization in 3D a-iCMs *(Scale bar: 5 μm)*.

### 2.3. 3D Hypertrophic Adipocytes Secretome Alters Cardiac Function of 2D a-iCMs

#### 2.3.1. Beating Kinetics

Due to the altered metabolic function of the myocardium, the obese heart is in an energetically disadvantaged position^82^. Since metabolic function is directly and critically related to the beating of CMs^83^, failing human CMs exhibit reduced contractility^84^. We recorded the spontaneous beating of the a-iCMs treated with 3D Adi/OAdi ConM and visualized the beating kinetics using vector heat maps to quantify alterations in the contractile dynamics of a-iCMs. The beating velocity of the 3D OAdi ConM treated a-iCMs was significantly decreased compared to 3D Adi ConM group (p=0.0312, Figure 3A), whereas beating frequency between groups was not significantly altered (Figure 3B). Beating confluency was substantially decreased in 3D OAdi ConM treated a-iCMs compared to 3D Adi ConM (Figure 3C). In supplementary studies, we once again included leptin low/high and adiponectin low/high groups to characterize the effect of major adipokines on the beating characteristics of a-iCMs (SFigure 5). The beating velocity of 3D OAdi ConM treated a-iCMs was significantly reduced compared to both adiponectin-containing groups (p=0.0106 and p=0.0070), in addition to the 3D Adi ConM group (SFigure 5A). Additionally, leptin-containing groups had a similar decrease with 3D OAdi group in terms of beating velocity compared to 3D Adi ConM and adiponectin-containing groups (SFigure 5A), indicating that leptin and adiponectin components released by adipocytes might be players in alterations in contractile function of a-iCMs in obesity. Beating frequencies remained relatively consistent across all groups (SFigure 5B), and neither adiponectin nor leptin supplementation resulted in significant differences in beating confluency (SFigure 5C), suggesting that while these adipokines influence contractile velocity, they may not substantially affect beat coordination or cell synchronization under the tested conditions.

**Figure 5:**
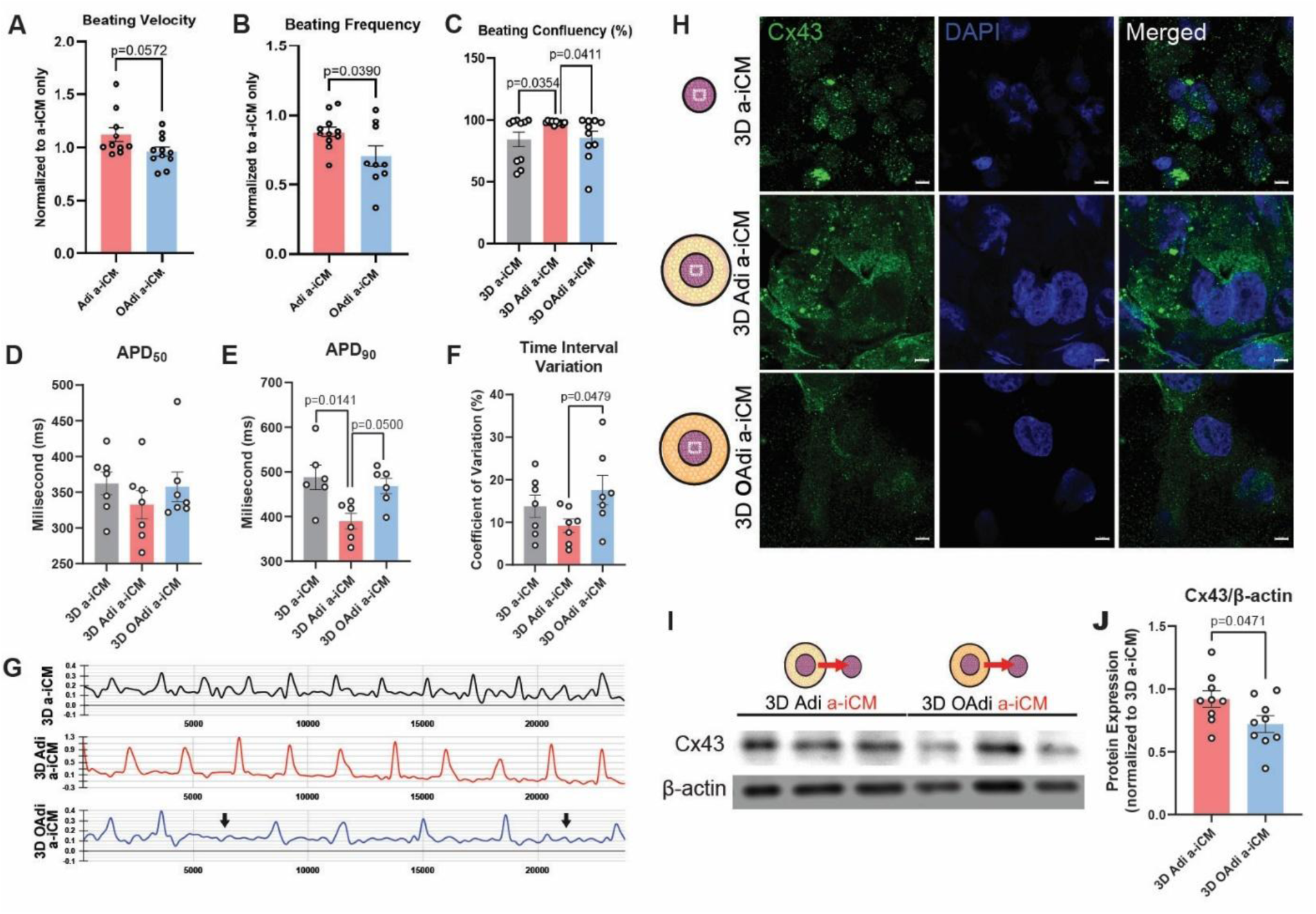
OAdi a-iCM constructs exhibit impaired contractile function, electrophysiological irregularities, and disrupted gap junction expression. A) beating velocity (n=11), B) beating frequency (n=9-11), and C) beating confluency (n=10) of a-iCMs cultured either alone, with 3D Adi, or 3D OAdi, D) APD50 (n=7), E), APD90 (n=6), F) time-to-peak variability (n=7), G) representative calcium transient waveforms H) Immunofluorescence staining for Cx43: green and DAPI: blue *(Scale bar: 5 μm)* I) Western blot analysis and J) quantification of Cx43 normalized to β-actin.

#### 2.3.2. Calcium Transient

Analysis of calcium transients is considered complementary to the beating kinetics of CMs, as calcium ions enter the cells during each beat and contribute to the electrical signal^85^. We recorded the calcium transient during spontaneous beating and quantified 50% (APD50, early repolarization), 90% decay times (APD90, late repolarization), and time to peak (TTP) variability. While APD50 and APD90 remained relatively consistent (Figure 3D-E), TTP variability increased substantially in the a-iCMs treated with 3D OAdi ConM compared to the B22 control (p=0.0687, Figure 3F), indicating potential disturbances in contraction timing or calcium handling in a-iCMs treated with 3D OAdi ConM. Additionally, we observed peak intensity increase in a-iCMs treated with 3D Adi ConM (Figure 3G). Taken together, results suggest that secreted factors from 3D OAdi negatively impact contractile function and calcium handling in a-iCMs while factors secreted by 3D Adi may enhance cardiac function of a-iCM. These findings indicate that lean adipocytes support metabolic and contractile function, while the detrimental paracrine effects arise from adipocyte dysfunction rather than adipocytes per se, underscoring the importance of adipocyte phenotype in modulating myocardial physiology.

### 2.4. 3D Hypertrophic Adipocytes Secretome Induces Intracellular Fat Accumulation in a-iCMs

Under physiological conditions, CMs store a small amount of intracellular triglycerides within lipid droplets, serving as an energy reserve to meet the high ATP demand of the heart, primarily through fatty acid oxidation. This finely tuned balance between lipid uptake, storage, and oxidation preserves cardiac metabolic flexibility^86^. However, in obesity, excess circulating FFA disrupts this balance, leading to ectopic fat accumulation within the myocardium^87^. Consequently, surplus lipids not only increase myocardial triglyceride content but also elevate levels of lipotoxic intermediates such as diacylglycerols and ceramides^88^, leading to increased mitochondrial and endoplasmic reticulum (ER) stress, and triggers proinflammatory and proapoptotic pathways^89^. Nile Red staining revealed minimal lipid accumulation in B22-treated (control) cells, a modest increase in cells exposed to 3D Adi ConM, and a major increase in intracellular lipid deposition in a-iCMs treated with 3D OAdi ConM (Figure 3H). Quantification confirmed significantly higher integrated lipid density per cell in the OAdi ConM group compared to both B22 (p=0.0044) and Adi ConM groups (p=0.0351, Figure 3I). These findings suggest that the 3D OAdi ConM, enriched in fatty acids, drives ectopic fat accumulation in monolayer a-iCMs, which may be driving impaired contractility, calcium handling, mitochondrial and glycolytic dysfunction, and metabolic inflexibility we reported (Figure 2-3).

### 2.5. Engineering and characterizing the 3D Fat-Myocardium Model

#### Cell Colocalization and Viability of the Model

After determining the direct paracrine effects of 3D Adi/OAdi ConM on a-iCMs, we constructed a fully 3D model (Figure 4A) to more accurately recapitulate the spatial and cellular interactions between adipocytes and CMs, enabling evaluation of both paracrine and juxtacrine mechanisms in a physiologically relevant microenvironment. The concentric circle model (Figure 4B) was designed to mimic the anatomical arrangement where EAT sits directly atop the myocardium, sharing the same microcirculation and enabling direct cell-cell and paracrine interactions. To colocalize the cells, 3D Adi and OAdi constructs were tagged with Orange CellTracker. a-iCMs were tagged with Deep Red CellTracker, incorporated into the GelMA-collagen hybrid bioink as explained in Section 4.4. and 3D bioprinted into the middle of the 3D Adi and OAdi constructs. 24h after a-iCM bioprinting, cell viability was assessed with Live/Dead staining. All 3D bioprinted groups (3D Adi/OAdi (SFigure 6A), and 3D a-iCM) had high viability (>93%) (Figure 4C, SFigure 6B). CellTracker-tagged adipocytes and a-iCMs were colocalized in the construct (Figure 4D-E). In parallel studies, constructs were stained using cell-specific markers, MLC2A (atrial CM specific, Figure 4F-G) and Nile Red (lipid specific, Figure 4F). Additionally, phalloidin localization in the 3D a-iCMs after 5 days in culture revealed well-organized cytoskeletal architecture and aligned CM, indicating a-iCMs established structural integrity after 3D bioprinting (Figure 4H).

### 2.6. Effect of 3D Hypertrophic Adipocytes on Beating Activity and Gap Junction Expression of 3D a-iCMs

#### 2.6.1. 3D Hypertrophic Adipocytes Alter Cardiac Function of 3D a-iCMs Beating Kinetics

Beyond paracrine signaling, the direct physical interaction between adipocytes and CMs plays a critical role in influencing cardiac function of CMs^40^. We captured the spontaneous beating of the 3D a-iCMs cultured with 3D Adi/OAdi cultures and visualized their kinetics using vector heat maps. The maps revealed that a-iCMs cultured with 3D Adi have tissue-like beating with high confluency (SFigure 7A), in contrast to a-iCMs cultured with 3D OAdi or alone. a-iCMs cultured with 3D OAdi exhibited a substantial decrease in beating velocity (Figure 5A) and a significant decrease in beating frequency compared to 3D Adi group (p=0.0390, Figure 5B). The significant decrease in beating velocity observed in a-iCM cocultured with 3D OAdi or a-iCM treated with 3D OAdi ConM aligns with prior findings in which rat CMs exposed to human white adipocyte-ConM exhibited an acute reduction in contraction amplitude^90^. In addition, a-iCMs cultured with 3D Adi had significantly higher beating confluency with minimal variability compared to a-iCMs cultured with 3D OAdi (p=0.0411) or a-iCM control (p=0.0354, Figure 5C). 3D OAdi a-iCMs demonstrated irregular, skipped beats as shown in Supplementary material: Movie 1.

**Figure 7.**
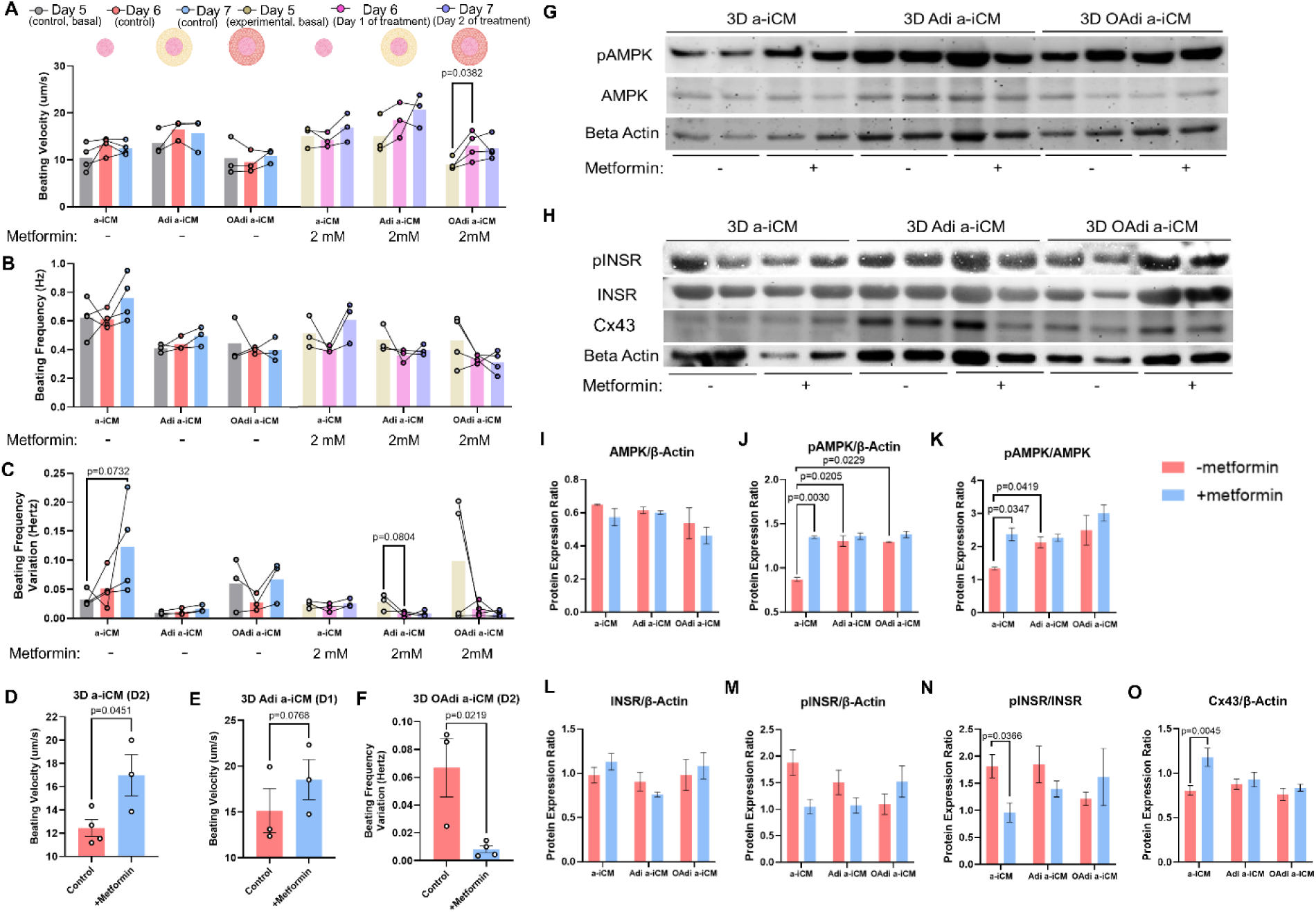
Metformin restores contractile function in OAdi-a-iCM constructs. A) Beating velocity (n=3-4) B) Beating frequency (n=3-4) and C) Beating frequency variation of 3D constructs treated with 2Mm metformin (n=3-4) D) Beating velocity comparison of -/+ metformin 3D a-iCM constructs (n=3-4) E) Beating velocity comparison of -/+ metformin 3D Adi a-iCM constructs (n=3) F) Beating frequency variation comparison of -/+ metformin 3D OAdi a-iCM constructs (n=3-4) G) AMPK, pAMPK and β-actin western blots. H) INSR, pINSR, and β-actin western blots. Western blot protein expression quantification of I) AMPK/β-actin (n=2), J) pAMPK/β-actin (n=2), K) pAMPK/AMPK (n=2), L) INSR/β-actin (n=4-10), M) pINSR/β-actin (n=4-10), N) pINSR/INSR (n=4-10), and O) Cx43/β-actin (n=4-10).

Compared to the ConM treatment, where a-iCMs exposed to 3D OAdi ConM showed reduced beating velocity without significant changes in beating frequency, the 3D coculture model resulted in more pronounced impairments. a-iCMs cocultured with 3D OAdi exhibited a marked decrease in beating velocity, a significant reduction in beating frequency, and significantly lower beating confluency compared to both 3D Adi and a-iCM-only controls (Figure 5A-C). These findings suggest that direct physical interaction with 3D OAdi may exacerbate contractile dysfunction of a-iCMs beyond the effects of secreted factors alone, emphasizing the importance of cell-cell and cell-matrix interactions in mimicking obesity-induced cardiac dysfunction.

##### Calcium Transient

In obesity, ectopic lipid accumulation in myocardium impairs mitochondria and the ER which disrupts calcium handling and promotes arrhythmias via delayed afterdepolarizations^23^. To investigate how these mechanisms manifest in our models, the calcium transient during spontaneous beating of 3D bioprinted a-iCMs, Adi a-iCMs, and OAdi a-iCMs were recorded. APD50, APD90, time-to-peak, and time interval variation were analyzed. APD50 remained relatively unaffected across models (Figure 5D), which was correlated to *in vivo* findings in ovine model, where short-term HFD exposure to ovine left atrial CMs had no significant impact on APD50^21^. We observed a marked increase in APD90 of a-iCMs cultured with 3D OAdi compared to a-iCMs cultured with 3D Adi (p=0.05, Figure 5E), suggesting impaired late-phase repolarization consistent with disrupted calcium reuptake dynamics. Interestingly, we observed a significant decrease in APD90 of a-iCMs cultured with 3D Adi compared to a-iCMs cultured alone (p=0.0141, Figure 5E). Although faster repolarization can, in some contexts, be pro-arrhythmic by increasing susceptibility to early afterdepolarizations^91^, the consistently high beating confluency, regular contraction patterns, and lack of arrhythmic features we observed in this group suggest otherwise. Taken together with our these findings and the fact that APD90 values for human atrial CMs typically range from 200 to 400 ms^92^, with the 3D Adi group averaging 389.81 ± 16.05 ms, we interpret this shortening as a sign of improved electrophysiological function with more efficient calcium reuptake dynamics or altered potassium channels.

Interbeat interval (IBI) variability, reflects the electrical stability and rhythmicity of CMs, and increased variability is often linked to arrhythmias^93,94^. Increase IBI variability could indicate disruptions in electrical conduction, and triggered activity, such as early or delayed afterdepolarizations, commonly seen in obese patients^95^. These abnormalities increase the risk of asynchronous contractions and reentry circuits, key mechanisms of arrhythmogenesis^96^. 3D OAdi a-iCMs showed significantly higher time interval variation compared to the Adi 3D a-iCM (p=0.0479, Figure 5F). Calcium transient waves of 3D a-iCMs and 3D Adi a-iCMs were regular, indicating consistent calcium cycling (Figure 5G).

3D Adi a-iCMs and control 3D a-iCMs exhibited regular and coordinated beating patterns. In contrast, 3D OAdi a-iCMs demonstrated irregular transients with skipped beats (indicated by arrows) with increased time interval variation, resembling atrial fibrillation, similar to the BF video analysis (Figure 5G, Supplementary material: Movie 1-2). This is further supported by the overlaid calcium transient peaks (SFigure 7B), where the 3D Adi a-iCM model exhibits sharper and more synchronized peaks compared to the broader and inconsistent peaks in the 3D OAdi a-iCM model. Correlated to our results, studies in obese sheep have demonstrated EAT expansion leads to conduction abnormalities and a higher risk of atrial arrhythmias^97^. However, AF rarely occurs naturally in animals and usually requires external induction, making it challenging to study its spontaneous onset and progression as in humans^98^. Therefore, our findings demonstrate that the 3D OAdi a-iCM model effectively mimics obesity-induced beating irregularity, offering a physiologically relevant platform to study AF-like phenotypes in the absence of external induction. These findings suggest that lean, 3D Adi created a supportive microenvironment that maintains/enhances a-iCMs calcium handling, rhythmicity, and synchronization, while the pathological effects of hypertrophic, 3D OAdi impair these critical functions.

#### 2.6.2. 3D Hypertrophic Adipocytes Alters Gap Junction Expression of a-iCM

Gap junction channels are essential for electrical conduction and synchronized contraction by enabling ion and small molecule exchange between myocardial cells^99^. Connexin43 (Cx43) is the predominant cardiac gap junction protein responsible for facilitating direct electrical and metabolic coupling between CMs, ensuring synchronous impulse propagation and coordinated contraction^100^. Remodeling, downregulation, or dephosphorylation of Cx43 has been strongly associated with impaired electrical conduction, increased arrhythmic susceptibility, and heart failure in both animal models^101^ and human cardiac disease^102^. Given the observed disturbances in calcium handling, rhythmicity, and excitation-contraction coupling in the 3D OAdi a-iCM model, we next investigated whether these functional impairments were linked to alterations in gap junctions.

Localization and comparison of Cx43 expression revealed differences between 3D Adi a-iCMs and 3D OAdi a-iCMs. Immunostaining shows that while 3D Adi a-iCMs display abundant, well-organized Cx43 at the intercalated disks, 3D OAdi a-iCMs exhibit visibly reduced and disorganized Cx43 distribution (Figure 5H). Cx43 protein expression obtained by western blot confirms this observation, with total Cx43 expression significantly lower in the 3D OAdi group compared to 3D Adi a-iCMs (p=0.0471, Figure 5I-J). The reduction in Cx43 abundance could compromise gap junctional communication, which likely contributed to the impaired calcium flux, disrupted rhythmicity, and excitation-contraction defects observed in the 3D OAdi a-iCM model. These findings highlight that hypertrophic adipocytes not only disrupt CM function through metabolic and paracrine effects but also impair electrical coupling by downregulating critical gap junction components. This provides important mechanistic insight into how obesity-driven adipose dysfunction may increase arrhythmic risk and promote cardiac dysfunction, underscoring the value of the 3D OAdi-a-iCM model for studying obesity-related cardiac pathology.

### 2.7. 3D Hypertrophic Adipocytes Decrease Insulin Receptor Phosphorylation and Glucose Uptake of a-iCMs

INSR are abundant in the heart and vasculature, regulating cardiac growth, survival, metabolism, and stress responses^103^. While insulin signaling adapts to metabolic changes to protect the heart from cardiotoxicity, its dysregulation in obesity and diabetes may contribute to myocardial dysfunction and heart failure progression^104^. In the atrium, insulin resistance induces atrial structural remodeling and disrupts intracellular calcium homeostasis, increasing susceptibility to arrhythmias^105^. It also induces mitochondrial dysfunction, ER stress and the unfolded protein response, oxidative stress, and inflammation^106^. To investigate the effects of hypertrophic adipocytes on INSR prevalence and activation, we assessed INSR and phosphorylated INSR (pINSR) expression in constructs. While INSR/β-actin expressions of 3D Adi a-iCMs and 3D OAdi a-iCMs remained relatively similar (Figure 6F), pINSR/β-actin expression of 3D Adi a-iCMs was substantially higher than 3D OAdi a-iCMs (Figure 6A, E). IF and western blot analysis revealed significantly decreased pINSR activation in 3D OAdi a-iCM compared to Adi a-iCMs (p=0.0122, Figure 6G). These findings indicate that while adipocyte hypertrophy does not significantly alter INSR expression, it impairs INSR phosphorylation in a-iCMs, potentially leading to insulin insensitivity, a hallmark of diabetes mellitus^107^.

**Figure 6:**
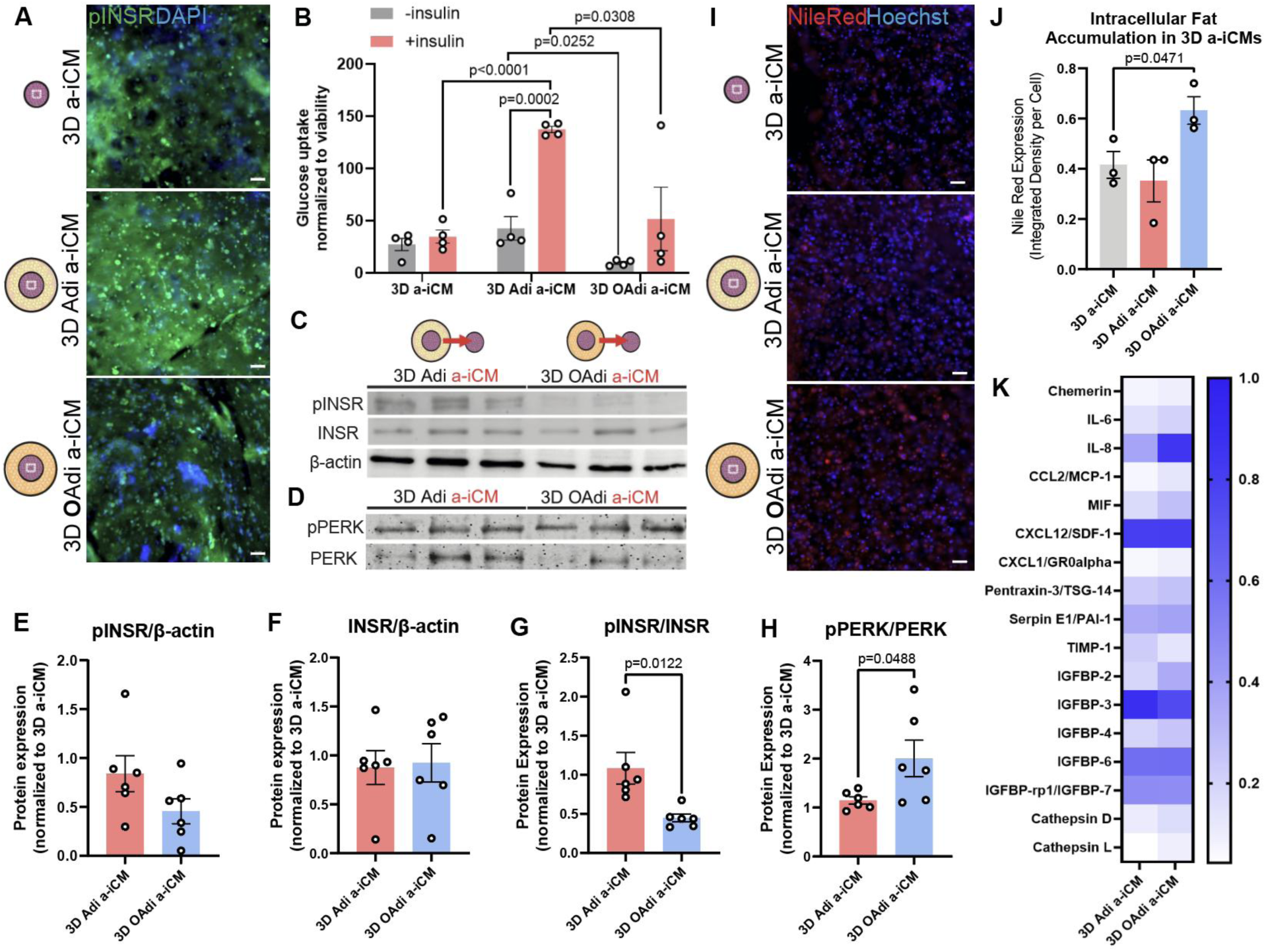
3D OAdi impairs glucose uptake and insulin signaling, and induces endoplasmic reticulum stress in a-iCMs. A) Representative immunofluorescence images of phosphorylated insulin receptor (green: pINSR) and nuclei (blue:DAPI) of the constructs *(scale bar=50 µm)*. B) Basal and insulin-dependent glucose uptake assay of 3D a-iCM portion of constructs (n=4) C) Western blot analysis of pINSR, INSR and D) pPERK, PERK in 3D a-iCMs cultured with Adi/OAdi. Quantification of E) pINSR/β-actin (n=6) F) INSR/β-actin (n=6) G) pINSR/INSR (n=6) H) pPERK/PERK (n=6) I) Representative lipid (red:Nile Red) and nuclear staining (blue:Hoechst33342) showing intracellular lipid accumulation in constructs *(scale bar = 50 µm)*. J) Quantification of Nile Red integrated intensity per cell (n =4). K) Cytokine analysis of 3D Adi/OAdi a-iCMs.

Reduced insulin receptor autophosphorylation is an early defect seen in the skeletal muscle cells of fat-fed, insulin-resistant rats^108^, and similarly, it was previously shown that elevated plasma FFA levels disrupted insulin signaling and glucose uptake in human skeletal muscle^109^. In the heart, studies in animal models showed that elevated circulating FFAs induced cardiac dysfunction and reduced myocardial glucose transporter expression^110^, and that chronic fatty acid exposure impaired both basal and stimulated glucose uptake^111^. In *in vitro*, palmitate treatment of human embryonic stem cell-derived CMs for 16 hours led to decreased insulin-stimulated glucose and fatty acid uptake^112^. Here, we compared basal and insulin-stimulated glucose uptake of 3D a-iCMs cultured with 3D Adi/OAdi for 5 days and observed that insulin significantly enhanced glucose uptake in 3D Adi a-iCMs (p=0.0002), whereas 3D OAdi a-iCMs showed no significant insulin responsiveness (Figure 6B). Basal glucose uptake was also significantly higher in 3D Adi a-iCMs compared to 3D OAdi a-iCMs (p=0.0252, Figure 6B), suggesting that obese adipocytes impair both basal and insulin-stimulated glucose uptake in a-iCMs. Here, we report that even without additional fatty acid supplementation to the CMs, HFD-fed adipocytes are sufficient to disrupt insulin signaling and impair glucose uptake in CMs. Under normal physiological conditions, lean adipocytes promote CM’s insulin sensitivity by secreting anti-inflammatory adipokines like adiponectin and omentin, which support glucose metabolism and reduce oxidative stress in the heart^8,35^. Consistent with this, our results showed that incorporating lean adipocytes into a-iCMs culture significantly enhanced insulin-stimulated glucose uptake compared to a-iCMs alone (p<0.0001, Figure 6B). Interestingly, a-iCMs cultured alone did not exhibit a significant increase in glucose uptake upon insulin stimulation, which may be due to their high basal reliance on glycolysis (Figure 2J) and limited insulin responsiveness in the absence of adipocyte-derived factors, an observation that warrants further investigation.

### 2.8. 3D Hypertrophic Adipocytes Increases Endoplasmic Reticulum Stress and Promote Intracellular Fat Accumulation in a-iCMs

The ER is a central organelle for protein folding, lipid synthesis, and calcium homeostasis in CMs^113^. Under physiological conditions, the ER ensures proper posttranslational folding and trafficking of membrane and secretory proteins^114^, while maintaining calcium signaling critical for excitation-contraction coupling^115^. However, when exposed to metabolic stressors such as excess FFA, the folding capacity of the ER becomes stressed^116^, leading to an accumulation of misfolded or unfolded proteins. In an attempt to regulate ER stress, cells activate the unfolded protein response (UPR) primarily through ER stress sensors such as protein kinase R-like ER kinase (PERK)^117^. While UPR initially promotes cellular adaptation by enhancing chaperone expression, reducing protein translation, and increasing ER-associated degradation, chronic or unresolved ER stress drives maladaptive responses, including oxidative stress, inflammation, calcium mishandling, and apoptosis, contributing to cardiac dysfunction. We hypothesized that the OAdi cultured a-iCMs, cultured in an environment with high FFAs and proinflammatory cytokines such as IL-6 and MIF, would have higher ER stress. Western blot analysis showed a significant increase in the phosphorylated PERK (pPERK)/PERK ratio in OAdi a-iCMs compared to both Adi a-iCMs (p=0.0488, Figure 6H), indicating that hypertrophic adipocyte co-culture increases ER stress sensor prevalence in a-iCMs. Consistent with the 3D OAdi ConM treated a-iCMs (Figure 3H-I), a-iCMs cocultured with 3D OAdi had significantly increased ectopic lipid accumulation compared to control 3D a-iCMs (p=0.0471, Figure 6I–J). In a clinical study, it was found that patients with impaired glucose tolerance and T2D exhibit significant myocardial triglyceride accumulation, known as cardiac steatosis, a condition considered a preclinical marker of diabetic cardiomyopathy^118^. This lipid overload occurs early in the disease course, before the onset of diabetes-related myocardial dysfunction, and is strongly associated with visceral adiposity rather than BMI or serum lipids^118^. In our study, by excluding exogenous lipids and exposing a-iCMs only to lipids derived from the adipose depot, we demonstrated that hypertrophic-adipocyte derived factors alone are sufficient to induce cardiac steatosis. This lipid buildup likely reflects enhanced FFA uptake and ectopic fat deposition within 3D a-iCMs cultured with 3D OAdi, which may exacerbate lipotoxic stress and contribute to ER stress, metabolic dysfunction, and impaired contractility.

### 2.9. Paracrine Signatures of Hypertrophic Adipocytes-a-iCM Model Reflect the Proinflammatory Phenotype of the Obese Heart

To assess the paracrine factors secreted by the engineered tissues, we analyzed the secretome profiles of 3D Adi a-iCM and 3D OAdi a-iCM constructs using a human cytokine array (Figure 6K). IL-6, IL-8, MIF and MCP-1/CCL2 were markedly increased in the 3D OAdi a-iCM secretome which are reported to be elevated in the circulation of obese individuals compared to lean subjects and is linked to low-grade inflammation, insulin resistance, atherosclerosis, and CVD^119,120,121,122^. Additionally, cathepsins were modestly upregulated in 3D OAdi a-iCM constructs, consistent with reports of increased cathepsin expression under conditions of cardiac stress, remodeling, and dysfunction^123^. Circulating cathepsin D levels are significantly increased in newly diagnosed type 2 diabetes, correlating with insulin resistance and early cardiac dysfunction^124^, while elevated cathepsin L levels are associated with increased cardiovascular mortality in older adults^125^. These findings suggest that the 3D OAdi a-iCM model recapitulates some key paracrine features of the obese heart, highlighting its relevance for studying obesity-associated cardiac dysfunction.

### 2.10. Metformin Mitigates Hypertrophic Adipocyte-Induced Impairments in a-iCMs

#### 2.10.1. Metformin Restores Contractile Function and Decreases Beating Frequency Variation of 3D OAdi a-iCMs

Animal models of CVD have shown that metformin improves cardiac function, primarily through activation of AMPK^126,127,128^. To assess the therapeutic potential of metformin in mitigating OAdi-induced dysfunction of a-iCMs, we treated the constructs with 2 mM metformin for 48 hours. Metformin significantly improved beating velocity in OAdi-a-iCM constructs after 24 hours compared to basal levels (p=0.0382, Figure 7A), suggesting enhanced contractile function. In 3D a-iCM constructs, metformin significantly increased beating velocity after 2 days of treatment compared to controls (no treatment) (p=0.0451, Figure 7D). In 3D Adi-a-iCM constructs, metformin led to a trend toward increased beating velocity 24h after treatment (p=0.0768, Figure 7E). Beating frequency across the construct groups (a-iCM, Adi-a-iCM, OAdi-a-iCM) remained relatively unchanged between untreated and metformin-treated conditions over 48 hours (Figure 7B). a-iCM constructs after 48 hours of treatment (p=0.0732) and Adi-a-iCM constructs after 24 hours of treatment showed a substantial decrease in beating frequency variation following metformin treatment (p=0.0804, Figure 7C). OAdi-a-iCM constructs exhibited higher basal beating frequency variation compared to other groups, which was markedly reduced following metformin treatment (Figure 7C). In 3D OAdi-a-iCM, metformin significantly reduced beating frequency variation after 2 days compared to control (no treatment) (p=0.0219, Figure 7F), a key metric associated with arrhythmias. This suggests that metformin may help mitigate arrhythmia-related dysfunction, aligning with previous reports showing its association with a reduced risk of new-onset atrial fibrillation in patients with obesity and T2D^129^.

#### 2.10.2. Metformin Activates AMPK Signaling in 3D a-iCMs

Metformin treatment led to notable activation of AMPK signaling across the constructs (Figure 7G). While total AMPK protein levels normalized to β-actin remained relatively unchanged with metformin in a-iCM, Adi-a-iCM, and OAdi-a-iCM constructs (Figure 7I), pAMPK/β-actin levels increased following treatment, particularly in a-iCMs (p = 0.0030, Figure 7J). A similar trend was observed in Adi-a-iCMs and OAdi-a-iCMs with a substantial increase in pAMPK/β-actin (Figure 7J). Importantly, the pAMPK/AMPK activation ratio also significantly increased in 3D a-iCM (p=0.0347) indicating that metformin enhances AMPK activity rather than its expression (Figure 7K). While a similar increase in AMPK activation was observed in Adi-a-iCMs and OAdi-a-iCMs, these changes did not reach statistical significance (Figure 7K). Significant effects of metformin on the contractile performance of a-iCMs (p=0.0451, Figure 7D) and the reduction in beating frequency variation in OAdi-a-iCMs (p=0.0219, Figure 7F) may be mediated through AMPK pathway activation. Interestingly, basal pAMPK/β-actin of 3D a-iCMs was significantly lower than a-iCMs in cocultures (p=0.0205, p=0.0229, Figure 7J). AMPK activation of a-iCMs cultured with 3D Adi was significantly higher than a-iCM cultured alone (p=0.0419, Figure 7K). This finding could suggest that the active metabolic crosstalk between adipocytes and a-iCMs, likely driven by adipokine signaling (particularly adiponectin^130^). These results underscore the ability of adipocytes, both lean and hypertrophic, to modulate AMPK signaling and CM metabolism, and metformin to markedly increase AMPK activity in 3D constructs.

#### 2.10.3. Metformin Alters Insulin Signaling and Gap Junction Expression of 3D a-iCMs

In liver cells, metformin has been shown to increase insulin receptor activation, particularly with IRS-2 signaling^131^, while in skeletal muscle cells, it was reported to increase insulin-stimulated tyrosine phosphorylation of the insulin receptor and IRS-1^132^. After metformin treatment, INSR expression remained relatively consistent across groups (Figure 7L); however, pINSR/β-actin levels and INSR activation levels in OAdi a-iCM showed a marked increase (Figure 7M-N). Interestingly, pINSR/β-actin levels and INSR activation in 3D a-iCMs were significantly lower (p=0.0366, Figure 7N). This could be due to AMPK activation reducing the reliance on INSR signaling, as previous studies have shown that AMPK activation can independently promote GLUT4 translocation to the cell membrane through insulin-independent pathways^47^. However, this mechanism in CMs warrants further investigation.

It was previously reported that metformin can restore connexin40 and Cx43 concentrations in a murine atrial fibrillation model^133^. Given the improvement in cardiac function observed with metformin treatment, we analyzed the expression of Cx43, which was previously shown to be decreased in our studies (Figure 5I-J). Metformin treatment significantly increased Cx43 expression in a-iCM constructs compared to untreated controls (p=0.0045), however, no significant changes in Cx43 expression were observed in Adi-a-iCM or OAdi-a-iCM constructs following metformin treatment (Figure 7O).

Taken together, metformin treatment improved contractile function and reduced beating frequency variation in 3D OAdi-a-iCM constructs, with a significant increase in beating velocity and a marked decrease in frequency variation after 48 hours. These functional benefits were accompanied by a modest increase in AMPK and INSR activation. Collectively, these findings highlight metformin’s ability to mitigate hypertrophic adipocyte-induced impairments in cardiomyocyte function, possibly through AMPK activation, modulation of insulin signaling, and preservation of gap junction integrity. Additionally, significant AMPK activation and increased Cx43 expression were also observed in a-iCM cultures alone, indicating that metformin’s functional and metabolic benefits are evident even in the absence of adipocyte co-culture.

## 3. Conclusions and Future Works

Despite the recognized importance of fatty acid component in cardiac tissue culture^1^, there is a notable absence of human-derived 3D models that incorporate both cardiac and adipose tissue elements. This gap in research limits our ability to fully understand the interactions between adipose tissue and myocardium in various (patho)physiological contexts, highlighting the need for the development of more comprehensive *in vitro* models. Here, we engineered a human-derived, 3D bioprinted fat-myocardium model designed to replicate the anatomical interface between EAT and atrial myocardium in obesity. For this, we first engineered 3D bioprinted hypertrophic adipocyte constructs that recapitulated key obesity hallmarks, including enlarged lipid droplets, altered cytoskeleton, increased lipolysis, proinflammatory cytokine release, impaired insulin signaling, and glucose uptake. We then investigated paracrine effects of hypertrophic adipocytes on a-iCMs using the secretome of 3D Adi/OAdi, which induced metabolic inflexibility in a-iCMs, marked by reduced mitochondrial respiration, glycolytic capacity, and ATP production, as well as contractile and calcium handling defects. Then, we integrated 3D a-iCMs with the adipocyte constructs to create a concentric, spatially defined coculture system. This model allowed for controlled paracrine and physical interactions while preserving the ability to independently retrieve and analyze each tissue component without reliance on single-cell sequencing or advanced sorting techniques. The 3D coculture revealed that direct interaction with hypertrophic adipocytes further impaired a-iCM beating kinetics, reduced beating confluency, and downregulated the gap junction protein Cx43, resembling key features of obesity-induced atrial dysfunction. Metformin treatment improved contractile function and reduced beating frequency variation in 3D OAdi-a-iCM constructs, with a significant increase in beating velocity and a marked decrease in frequency variation as well as a modest increase in AMPK and INSR activation. Overall, human-derived 3D bioprinted fat-myocardium model successfully recapitulates key structural, metabolic, and electrophysiological features of obesity-induced atrial dysfunction and demonstrates metformin’s ability to mitigate contractile and metabolic function of a-iCMs cultured with hypertrophic adipocytes.

Future iterations of our model could benefit from the integration of iPSC-derived adipocytes, which offer a renewable and scalable alternative to primary ADSCs. Unlike ADSCs, whose adipogenic potential declines with passaging, iPSCs enable consistent adipocyte generation and hold promise for modeling EAT-myocardium interactions and offer high-throughput drug screening. Additionally, enhancing the maturity of hiPSC-derived CMs remains a critical avenue for improvement. Although these cells provide chamber specificity, they retain fetal-like structural and metabolic characteristics that may limit their fidelity in modeling adult atrial pathophysiology^75^. Efforts to promote iCM maturation, such as mechanical/electrical stimulation^134,135^ or tailored metabolic maturation media^75^, could improve their translational relevance.

While adipocytes in the current study were derived from non-cardiac depots (e.g., abdominal fat), future studies incorporating human EAT-derived stem cells could more precisely model depot-specific adipocyte behavior, further improving the physiological accuracy of the platform. Furthermore, in contrast to our study, which primarily investigated the effects of adipocytes on cardiomyocytes, future work could investigate how CM-derived factors, cardiokines, directly influence adipocyte function, including lipid metabolism, adipogenesis, and inflammatory signaling^136^.

This work provides mechanistic insights into adipocyte dysfunction-induced arrhythmia, mitochondrial impairment, and disrupted insulin signaling, and establishes a versatile platform for investigating obesity-induced CVD. Beyond mechanistic insights, the model holds translational potential for evaluating therapeutic agents, as demonstrated by metformin screening, and for guiding personalized treatment strategies for obesity-related atrial dysfunction.

## 4. Experimental Section

### 4.1. Human ADSC culture and 3D bioprinting

Primary ADSCs were isolated from the axilla, mid back, flanks, and central abdomen of patients as described previously^137^. Isolation and characterization of ADSCs were carried out at Indiana University School of Medicine by Dr. Nakshatri’s group, and the procedures to collect human adipose tissue were approved by the IU School of Medicine Institutional Review Board. Details on the characteristics of the ADSCs used in this study, including donor BMI, gender, and other relevant information, are provided in Supplementary Table 1.

GelMA hydrogels were synthesized utilizing an established protocol^138^, Human Skin Collagen type 1 (10 mg/ml, Humabiologics) and photoinitiator (2-hydroxy-4-(2-hydroxyethoxy)-2-methylpropiophenone, Sigma) were purchased. Degree of methacrylation was quantified using NMR, and GelMA lysine methylene area and gelatin lysine methylene area were calculated using MestreNova. The degree of methacrylation of synthesized GelMA hydrogels was determined as 54% (SFigure 8) using the equation we reported previously^139^. Mechanical properties of the bioink were characterized utilizing the frequency sweep test and nanoindenter, as we reported before^44^. GelMA-Collagen type 1 bioink had Young’s modulus of 2.0 ± 0.8 kPa^44^, which is within the reported value for both myocardium^140^ and adipose tissue^141^.

ADSCs were maintained in ADSC Growth Medium and trypsinized at ∼80% confluency. Collagen type I (1 mg/mL, final concentration) was combined with 5 million/mL ADSCs and mixed with photoinitiator (0.025%, final concentration) and GelMA (10%, final concentration). Bioink was loaded into cartridges and extruded through 22G nozzles onto sterilized, charged glass slides (Globe Scientific Inc.) using the CELLINK BioX6 bioprinter and a custom-generated G-code. Constructs were printed in a hollow cylindrical shape (8 mm outer diameter, 4 mm inner diameter, 0.3 mm height) following the parameters outlined in Supplementary Table 2, and subsequently, constructs were photocrosslinked for 30 seconds under UV light (6.9 W/cm²) using a UV lamp (Lumen Dynamics). Three days after bioprinting, ADSC constructs were differentiated using adipogenic induction media (Lonza) as explained previously^142^ and lipid droplet accumulation was validated by Nile Red (Invitrogen) staining. After the ADSC to adipocyte differentiation protocol (SFigure 3A), the intracellular accumulation of lipid droplets in adipocytes was validated using Nile Red staining (SFigure 1A) and BF imaging (SFigure 1B). HFD was recapitulated using DMEM/F12 1%FBS supplemented with 750 μM PA pre-conjugated to BSA (Cayman) to obtain hypertrophic adipocytes as described before^16^. Vehicle media supplemented with 80 μM of fatty acid-free BSA (Sigma Aldrich, USA) was added to the control group. 3D Adi/OAdi were incubated in DMEM/F12 1% FBS for 6 days with media changes done every 2 days, and ConM were collected for further analysis.

### 4.2. Characterization of 3D hypertrophic adipocytes

24 hours after fatty acid treatment, cells were stained with Nile Red to visualize intracellular lipid droplets, and the droplet sizes of 3D Adi and OAdi were visualized using fluorescence microscopy. TRITC-labelled F-actin (Invitrogen) immunostaining was conducted to visualize the cytoskeletal changes during HFD. Basal lipolysis was quantified by incubating the scaffolds in DMEM/F12 for 2 days at 37°C. The conditioned media were collected, and the Glycerol Glo™ Assay kit (Promega, WI, USA) was utilized to quantify basal lipolysis rate following the manufacturer’s protocol using a plate reader (SPARK plate reader, TECAN).

To quantify glucose uptake, 3D Adi/OAdi were washed and starved for 3 h (no FBS/no glucose). Then, 10 μM human insulin was added to the constructs for 1h. 1 mM 2DG in PBS was added to each well and incubated for 10 minutes at room temperature. Glucose uptake by 3D Adi and OAdi was assessed using the Glucose Uptake-Glo™ Assay Kit (Promega, WI, USA), and luminescence was recorded with a plate reader according to the manufacturer’s instructions. Luminescence values of 2DG uptake were normalized to cell viability, which was quantified using the CellTiter-Glo® Luminescent Cell Viability Assay (Promega, WI, USA).

The relative cytokine content of 3D Adi/OAdi constructs was determined using the human XL cytokine array (R&D Systems, Cat. No: ARY028) and human adipokine array (R&D Systems, Cat. No: ARY024) kits following manufacturer’s instructions. Briefly, cytokine array membranes were blocked with an array buffer for 1 hour at room temperature, washed, and incubated overnight at 4 °C in an equal volume of the conditioned media. The membranes were washed and then incubated with the biotinylated antibody cocktail solution for 1 hour, followed by incubation in streptavidin-horseradish peroxidase (HRP) for 30 minutes and the chemiluminescent reagent for 1 minute. The membranes were then exposed to X-ray for 15 minutes using a biomolecular imager (ImageQuant LAS4000, GE Healthcare). Relative cytokine levels were determined by quantifying the dot intensity using ImageJ. Commercial ELISA kits were used to quantify human adiponectin (KHP0041; Invitrogen, USA) and human leptin (KAC2281; Invitrogen, USA) following the manufacturer’s protocols. Insulin sensitivity was assessed by western blot quantifying INSR. Adipocyte constructs were snap-frozen and crushed using liquid nitrogen. Then, the resulting powder was incubated with RIPA buffer with 1% proteinase inhibitor cocktail at 4°C for 30 minutes. The protein concentration of each construct was quantified via bicinchoninic acid (BCA) assay (Thermo Fischer), and equal amounts of protein were separated by 10% SDS-PAGE and transferred to the blotting membrane. After exposure for 1-4 minutes, the pixel density of each protein band was quantified using ImageJ.

### 4.3. Differentiation of hiPSC-derived atrial cardiomyocytes

SeVA1016 (reprogrammed from human skin fibroblasts)^143^ hiPSCs (RRID: CVCL_UK18) were differentiated into a-iCM and v-iCM (for comparison) by adapting previously reported two-week small molecule-based treatment protocols (SFigure 9A)^144,145^. For a-iCM differentiation, once hiPSCs reached 80-95% confluency, cells were treated with CM (−) (RPMI Medium 1640, Corning) supplemented with B27 without insulin (2%, Gibco), beta-mercaptoethanol (β-ME) (final concentration of 0.1 mM, Promega) containing 5 μM CHIR99021 (Stemcell Technologies) for 48 hours. On Day 2, the medium was replaced with Cardio Differentiation Medium supplemented with 5 μM IWP-4 (Stemcell Technologies). On Day 3, 1 μM retinoic acid (Sigma-Aldrich) was added without media aspiration, followed by a fresh medium change with 1 μM retinoic acid on Day 4. On Day 6, the medium was replaced with CM (−). Starting from Day 8, cells were maintained in fresh CM (+) (RPMI Medium 1640 supplemented with B27 with insulin (2%, Gibco), β-ME (final concentration of 0.1 mM), with media changes every 3 days. Beating of a-iCMs and v-iCMs was observed by Day 15 of differentiation. On Day 30, differentiation efficacy and chamber specificity of the a-iCMs were assessed by immunostaining for sarcomeric alpha-actinin, troponin T (TnT), and MLC2A markers, applying concentrations as recommended by the manufacturer (Supporting Information: Table 3), and following the immunostaining protocol outlined in our previous work^44^. To validate MLC2A expression, we employed flow cytometry, using a previously established protocol^146^. IF results showed the striated cytoskeletal structures (SFigure 10A) and aligned Z discs (SFigure 10B). We characterized the chamber specificity of iCM cultures, using immunostaining (SFigure 9B), flow cytometry (SFigure 9C), and beating analysis (SFigure 9D-E). IF analysis revealed high expression of atrial cardiac marker MLC2A in a-iCM and ventricular marker MLC2V in v-iCM cultures (SFigure 9B). Video analysis of lateral displacement of spontaneous beating chamber-specific iCMs was generated using an in-house MATLAB code with the block-matching algorithm, as we previously described^147^. After differentiation, we analyzed the differentiation efficacy of iCMs by immunofluorescence and flow cytometry^148^, and the functionality of iCMs by analyzing beating and electrophysiological properties utilizing BF video analysis^44^. Consistent with the literature^144^, a-iCMs had significantly higher beating frequency than v-iCMs (SFigure 9E).

### 4.4. Characterizing the metabolic effects of hypertrophic adipocytes on a-iCM

Seahorse XF96 extracellular flux analyzer was used to assess mitochondrial function as we described before^149^. Briefly, a-iCMs were seeded at a density of 100,000 cells per Matrigel-coated well in Seahorse XF96 Microplates five days before the assay. 48 hours prior to the assay, 3D Adi/OAdi ConM was supplemented with 2% B22 with insulin and applied to monolayer a-iCMs. One hour prior to the assay, the culture medium was replaced with Agilent Seahorse XF RPMI Basal Medium supplemented with 2 mM glutamine, 10 mM glucose (no glucose for Glycolysis Stress Test), and 1 mM sodium pyruvate. For the Mito Stress test, the Seahorse XF Cell Mito Stress Test Kit (Agilent) was used, with inhibitors prepared at the following concentrations: oligomycin (2.5 μM), FCCP (2 μM), and a combination of rotenone and antimycin A (Rot/AA, 2.5 μM). For the Glycolysis Stress test, Seahorse XF Glycolysis Stress Test Kit (Agilent) was used, with inhibitors prepared at the following concentrations: glucose (10 mM), oligomycin (1 μM), and 2-DG (50 mM). For Glycolytic Rate test, Seahorse XF Glycolytic Rate Assay Kit (Agilent) was used, with inhibitors prepared at the following concentrations: Rot/AA (0.5 μM) and 2-DG (50 mM). The number of a-iCMs per well was quantified using Hoechst 33342 (Thermo Scientific, 8 µM) after a 30minute incubation at 37°C, followed by image analysis with ImageJ. OCR, ECAR, and PER were then normalized to the cell count. Baseline OCR was determined as the average of measurements taken from point 1 to 3, Basal Respiration, Maximal Respiration, Spare Respiratory Capacity, Coupling Efficiency, ATP Production-Coupled Respiration, MitoATP, Basal Glycolysis, Compensatory Glycolysis, GlycoATP, and % Proton Efflux Rate from Glycolysis were analyzed on Seahorse Analytics. Glycolytic Capacity (ECAR after oligomycin injection – ECAR before glucose injection), % Glycolytic Reserve ((Glycolytic Capacity – Basal Glycolysis) / Glycolytic Capacity) × 100), and Non-Glycolytic Acidification (ECAR before glucose injection) were derived from the raw data. These values were subsequently graphed and analyzed using GraphPad Prism. In supplementary experiments, additional treatment groups, Adiponectin (Human Recombinant Adiponectin, Abcam) (low (8 ng/ml)/high (12 ng/ml)), Leptin (Human Recombinant Leptin, VWR) (low (3000 pg/ml)/high (5000 pg/ml)), and PA (BSA-conjugated palmitic acid, Cayman) (low (250 μM)/high (500 μM)), were included, and a-iCM treatments were conducted using the same time frame. Adiponectin and leptin concentrations were selected based on trends observed in ELISA results. The 3D Adi group was characterized by high adiponectin and low leptin levels, whereas the 3D OAdi group exhibited low adiponectin and high leptin levels (Figure 1K).

### 4.5. Characterization of paracrine effects of hypertrophic adipocyte on a-iCM

To analyze the paracrine effects of hypertrophic adipocyte on a-iCM’s contractile function, ConM treatment detailed in 4.4. was repeated. After 48 hours, 3D Adi and OAdi ConM treated a-iCMs were compared in terms of beating velocity, frequency, and confluency using a video analysis generated using an in-house MATLAB code with the block-matching algorithm, as we previously described^147^. Then, electrophysiological changes induced by hypertrophic adipocytes were further analyzed by characterizing the calcium (Ca²⁺) transient of a-iCMs. Cells were washed with PBS, and the medium was replaced with a Ca²⁺-sensitive Fluo-4 AM (Life Technologies) solution, following the manufacturer’s instructions. Real-time beating videos of 3D a-iCMs were recorded using a fluorescence microscope equipped with a Hamamatsu C11440 digital camera, with a 30 ms exposure time for 20 seconds. The rate of Ca²⁺ release (time to peak Ca²⁺ transient amplitude) and action potential duration at 50% (APD50) and 90% (APD90) of the amplitude were derived from intensity versus time plots. Additionally, intracellular fat accumulation in a-iCMs treated with 3D Adi/OAdi ConM was characterized by Nile Red staining and quantified in ImageJ.

### 4.6. Engineering of hypertrophic adipocyte a-iCM co-culture model

To visualize cell colocalization and spatial distribution across tissue layers, ADSC-adipocytes were tagged with CellTracker orange CMTMR (1 µM, Life Technologies). At Days 30-35 of the differentiation, a-iCMs were tagged with CellTracker far red (1 µM, Life Technologies) and detached using trypsin-EDTA supplemented with 0.25 U/mL Liberase TH (Roche). Collagen type I was combined with a-iCM pellet and mixed with photoinitiator and GelMA. 90 million/mL a-iCM bioink was loaded into cartridges and extruded through 20G nozzles using the droplet bioprinting setting on the CELLINK BioX6 Bioprinter. The media was removed from the wells containing 3D bioprinted ADSC-derived lean and obese adipocytes prior to the second round of bioprinting. The bioink was bioprinted as droplets into the core of the 3D Adi/OAdi constructs and subsequently photocrosslinked for 30 seconds under UV light (6.9 W/cm²) using a UV lamp. In parallel, 3D a-iCM-only control groups were bioprinted onto sterilized charged glass slides (Globe Scientific Inc.) as controls using the same a-iCM bioink. All constructs were incubated in tailored coculture media with 5:1 RPMI Medium 1640:DMEM/F12 supplemented with 2% B27 with insulin. 24 hours after bioprinting, the tile images were taken with a fluorescence microscope (Zeiss, Hamamatsu ORCA flash 4.0) and stitched together using Zen Software.

### 4.7. Characterization of the hypertrophic adipocyte a-iCM co-culture model

24 hours after the a-iCM bioprinting, the viability of the 3D Adi/OAdi-a-iCM models was analyzed utilizing Live-Dead assay (Life Technologies) following the manufacturer’s instructions, and live/dead cells were counted using ImageJ, and %viability was calculated as reported before^44^. Constructs were stained using Nile Red, TnT, and MLC2A to colocalize adipocytes and a-iCM in the constructs. After five days in culture, a-iCMs in both cultures were compared in terms of beating velocity, frequency, and confluency using BF beating video analysis as well as Ca²⁺ transient, as detailed in section 4.5. Cx43, INSR, pINSR, PERK, and pPERK levels were compared between groups using immunostaining and western blot analysis. For Western blot analysis, the a-iCM and adipocyte components of the constructs were carefully separated on Day 5, snap-frozen immediately, and stored at -80°C. For immunostaining, constructs were washed and fixed with 4% PFA, and immunostaining protocol was applied as explained above. Glucose uptake of a-iCMs cultured with 3D Adi/OAdi was conducted as detailed in Section 4.2. Intracellular fat accumulation in 3D a-iCMs was characterized by Nile Red staining and quantified in ImageJ.

### 4.8. Metformin treatment of hypertrophic adipocyte a-iCM co-culture model

Five days after a-iCM bioprinting, 3D a-iCM (control), and Adi/OAdi-a-iCM were treated with 2 mM metformin (Sigma-Aldrich, MO, USA) for 48 hours. The treatment media was replaced after 24 hours to maintain drug activity and nutrient availability. Following treatment, constructs underwent the same characterization protocol described in section 4.5. Briefly, beating kinetics, and calcium transients were characterized as detailed in section 4.5. To examine molecular changes, protein expression levels of Cx43, INSR, pINSR, pAMPK, and AMPK were analyzed by western blotting. For western blots, constructs were disassembled to separate the a-iCM and adipocyte components, snap-frozen, lysed, and analyzed as described in sections 4.2. and 4.7.

## Acknowledgements

This work was supported by NIH Award #R01 CA275423-01A1 and NSF EFMA-2422333. We thank Nakshatri lab for providing human adipose-derived stem/stromal cells. We acknowledge the Biological Screening and Assay Development Core at the University of Notre Dame for their technical support with the Seahorse metabolic analyzer. We acknowledge the guidance of J. Yang (biochemical assays), G. Basara (3D bioprinting), A. Hansrisuk (NMR analysis), S. G. Ozcebe (Seahorse assays), and A. T. Huerta (flow cytometry). We also thank J. Yang, L. Hawthorne, G. Ronan, M. Zarodniuk, and F. Ketchum for their input on the manuscript.

